# DRGSCROLL: Achieving Full Side-Chain Flexibility in Docking Simulations through Genetic Algorithm Framework

**DOI:** 10.64898/2025.12.04.692419

**Authors:** Ehsan Sayyah, Muhammet Eren Ulug, Huseyin Tunc, Serdar Durdağı

## Abstract

Structure-based docking often assumes a rigid receptor, obscuring induced-fit side-chain rearrangements that govern affinity and selectivity. We introduce DRGSCROLL, an open access docking platform and web server that jointly optimizes ligand pose and continuous receptor side-chain χ angles within a genetic-algorithm (GA) optimizer. DRGSCROLL seeds broad χ-angle populations, evaluates candidates with a dual-objective fitness that rewards low interaction energy profile while penalizing steric clashes, and uses per-residue crossover plus stochastic mutation to maintain physically realistic χ sets. To favor exploration over premature local minima, per-iteration minimization is deliberately omitted; final poses are selected by clash-free filtering and RMSD-based clustering. Across PDBbind protein-ligand complexes, DRGSCROLL showed generation-wise decreases in clash counts and improved docking scores, indicating convergence to sterically viable, low-energy pocket conformations rarely accessed by rigid-receptor protocols. Relative to known publicly available and commercial docking programs such as Vina, Glide, and IFD; DRGSCROLL sampled more favorable energy distributions with lower median scores, consistent with superior induced-fit capture. In prospective virtual-screening evaluations, DRGSCROLL enhanced actives versus inactives discrimination for RET kinase and PARP1 targets, while maintaining balanced precision–recall trade-offs. Additionally, DRGSCROLL’s endurance in differentiating actives from property-matched decoys was validated by benchmarking against a decoy database (Directory of Useful Decoys, DUDE), which showed improved AUC, recall, and enrichment performance in comparison to Vina GPU 2.1 under the same screening settings. By embedding continuous side-chain flexibility directly into the search, rather than relying on post hoc rotamer tweaks or heavy local minimization, DRGSCROLL addresses key combinatorial and feasibility bottlenecks in flexible docking. The method provides a scalable, physics-grounded route to adaptive receptor-ligand modeling that improves pose accuracy and early enrichment for flexible targets, from fragment screening to triage of synthetically ready or AI-generated small molecule libraries. Availability: platform page, https://www.drgscroll.com/; academic/nonprofit web server, https://drgscroll.bau.edu.tr/

## 1. Introduction

Structure-based drug discovery (SBDD) has become a cornerstone of modern medicinal chemistry, enabling the rational design and optimization of therapeutic agents across diverse therapeutic areas ^1–3^. Among the computational strategies available, molecular docking remains central, aiming to predict the binding modes and affinities of small molecules for protein targets. Docking serves as both a hypothesis generator and a virtual screening filter, allowing millions of compounds to be prioritized for experimental testing at a fraction of the cost and time of conventional methods ^2^. Its success depends critically on two interdependent factors: (i) accurate sampling of plausible ligand poses within the binding site, and (ii) scoring functions capable of approximating binding free energies ^4,5^. Despite decades of refinement, the accuracy and robustness of docking remain fundamentally constrained by the treatment of protein flexibility.

Proteins are intrinsically dynamic macromolecules whose conformational plasticity is essential for ligand recognition, allosteric regulation, and catalysis. Among these structural adjustments, side-chain χ-angle rotations can dramatically reshape the steric, electrostatic, and hydrogen-bonding profile of the binding pocket ^4,5^. Neglecting such plasticity and treating receptors as rigid scaffolds often leads to misplaced ligands, distorted scoring outcomes, and misranking of active compounds ^1–3^. This limitation is especially acute for flexible targets such as kinases, ion channels, and enzymes with mobile loops, where induced fit is a dominant determinant of selectivity and potency.

Several strategies have been proposed to mitigate this problem. Induced-fit docking (IFD) protocols attempt to incorporate receptor relaxation by combining initial docking with rotamer sampling and local minimization of the binding site ^6^. Post-docking refinement methods, such as FiberDock, use normal-mode analysis and side-chain optimization, but these adjustments are localized and do not capture broader conformational diversity ^7^. FlexAID explicitly incorporates side-chain flexibility with soft scoring to tolerate non-native structures ^8^, while AutoDockFR allows user-specified flexible side chains ^9^. GOLD applies a genetic algorithm (GA) to efficiently explore ligand flexibility and is widely used in industry ^10^. However, GOLD treats the receptor largely as rigid or permits only limited side-chain sampling via predefined rotamers, thus missing continuous χ-angle plasticity. Despite their utility, these approaches are all hampered by rotamer approximations, limited side-chain exploration, or minimization-heavy refinements that compromise accuracy and scalability.

Other strategies distribute flexibility over multiple receptor conformations. Ensemble docking extends the traditional paradigm by incorporating a set of receptor conformations—obtained from crystallographic structures, NMR ensembles, or molecular dynamics trajectories—to account for the intrinsic plasticity of proteins. This strategy enables ligands to explore a wider range of binding-site geometries, often revealing alternative binding modes that single-structure docking would miss. Despite its conceptual elegance, ensemble docking faces practical limitations. As the number of receptor conformations increases, so does the computational cost and the likelihood of identifying spurious poses in conformations that are not functionally relevant. Recent studies have attempted to mitigate these issues by clustering molecular dynamics (MD)-derived snapshots or weighting conformers according to energetic or experimental criteria, highlighting a trade-off between conformational diversity and predictive precision. ^11,12^. MD-coupled workflows capture receptor plasticity with atomistic realism ^13^, yet their prohibitive computational expense limits applicability to large-scale screening campaigns. Ultimately, the field remains hindered by two persistent challenges: (i) the combinatorial explosion of χ-angles, which renders exhaustive conformational search infeasible ^4,11,12^, and (ii) scoring functions that penalize transient steric clashes or strained geometries arising naturally during conformational rearrangements ^5,13^. Together, these bottlenecks force most docking engines into oversimplified solutions that fail to capture the true energetic landscape of protein-ligand binding.

In parallel, machine learning (ML)-based docking and scoring has emerged as a transformative direction. Convolutional neural networks (CNNs) such as GNINA improve pose ranking and affinity prediction by learning directly from protein-ligand complex data ^14^. Geometric deep learning approaches, including EquiBind ^15^ and DiffDock ^16^, leverage graph neural networks (GNNs) and diffusion generative models to predict binding poses without traditional search heuristics. These innovations promise substantial acceleration of docking pipelines and the discovery of novel chemotypes overlooked by classical scoring. However, limitations remain: performance is strongly dependent on biased training datasets, generalization to novel or cryptic binding sites is uncertain, and predictions are often opaque. Critically, most ML models assume rigid receptor geometries and do not explicitly model side-chain χ-angle flexibility.

Thus, while ML-based approaches complement physics-based methods, they do not resolve the long-standing problem of receptor plasticity.

Here, we introduce DRGSCROLL (*Durdağı Research Group’s Side-Chain ROtations for Ligand Landing*), a next-generation open access docking platform (https://www.drgscroll.com) and webserver (https://drgscroll.bau.edu.tr) designed to directly address this unmet need. To our knowledge, DRGSCROLL is the first docking engine to integrate continuous χ-angle optimization of receptor side chains within an evolutionary search framework. Unlike conventional methods restricted to rotamer libraries (e.g., GOLD, AutoDockFR, FlexAID) or minimization-heavy induced-fit protocols, DRGSCROLL implements a GA that simultaneously explores ligand poses and receptor conformations in a continuous search space. Its core innovations include: (i) initialization by random sampling of χ-angles across a broad conformational landscape, avoiding premature pruning; (ii) a dual-objective fitness function that balances docking scores with steric clash penalties, embedding physical feasibility directly into the GA search; (iii) per-residue crossover operators that transmit intact χ-angle sets, preventing unphysical hybrids and accelerating convergence; (iv) stochastic mutations to preserve population diversity; and (v) the deliberate omission of explicit energy minimization during iterations, reallocating computational effort toward expansive sampling rather than repetitive relaxations.

By embedding steric feasibility into the GA itself, DRGSCROLL effectively resolves the combinatorial explosion problem that has long undermined receptor-flexible docking. This enables a scalable, physics-grounded exploration of receptor plasticity that surpasses the rotamer-limited frameworks of GOLD, FlexAID, and AutoDockFR, and complements the speed of ML-based scoring with a rigorous treatment of induced fit.

Benchmarking against state-of-the-art docking engines demonstrates that DRGSCROLL achieves superior pose accuracy and ligand ranking across diverse PDBbind complexes. More importantly, in functional virtual screening campaigns targeting RET tyrosine kinase (TK) and PARP1 inhibitors obtained from the ChEMBL dataset (https://www.ebi.ac.uk/chembl/), together with DUDE-derived decoys for PARP1 (https://dude.docking.org/targets/parp1), DRGSCROLL outperformed rigid and lightly flexible protocols, delivering higher enrichment metrics (i.e., AUC, F1, Precision, Recall, Accuracy). Beyond benchmarking, the algorithm provides a generalizable framework for applications ranging from fragment-based drug discovery to screening AI-generated ligand libraries against highly flexible binding sites.

In summary, DRGSCROLL represents a paradigm shift in receptor-flexible docking. By combining continuous χ-angle optimization with evolutionary search, it overcomes long-standing bottlenecks of combinatorial explosion and steric infeasibility. This advance elevates docking from static or discretized receptor models toward truly adaptive receptor-ligand modeling, providing a robust, scalable, and generalizable tool at the interface of physics-based and ML-enhanced drug discovery.

## 2. Methods

We constructed protein-ligand complexes using a staged workflow that integrates molecular docking with torsional sampling and GA-driven optimization of receptor side-chain conformations in the binding pocket. Proteins were prepared by assigning ionization states, adding hydrogens, and performing restrained optimization; ligands were processed by enumerating relevant protonation/tautomeric states, generating conformers, and refining geometries. Docking with ADFR ^9^ produced an ensemble of poses, from which a provisional complex was selected based on interaction geometry and docking affinity. Residues proximal to the ligand were then subjected to side-chain sampling, and GA-based χ-angle optimization refined local packing and interaction energetics. This procedure systematically captures induced-fit effects and yields energetically favorable complexes suitable for downstream analysis of protein-ligand recognition.

### 2.1 Residue Selection and Sidechain Rotation

Residues within a 4 Å distance from the ligand were identified using the following distance criterion:

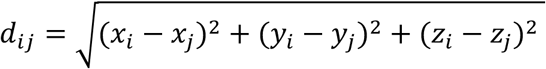

where *d_i,j_* is the Euclidean distance between the ligand atom *i* and the protein residue atom *j*. Residues satisfying *d_i,j_* ≤ 4.0 Å were selected for further conformational sampling. In order to exclude distant residues that contribute minimally to ligand affinity and to guarantee the inclusion of side chains that directly affect ligand binding, this cutoff was selected. Other filtering criteria were used when choosing residues for optimization. Proline (Pro) and glycine (Gly) residues were omitted because they lack rotatable side-chain torsions. A check was made to see if there was a disulfide bond present in cysteine (Cys) residues. To maintain the integrity of the disulfide bond, a Cys residue was specifically barred from further rotation if another sulfur atom was discovered within 2.05 Å of the sulfur atom. The following was the formulation of the Cys residue selection criterion:

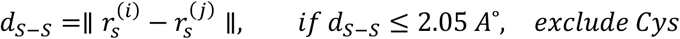

where 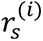 and 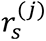 are the coordinates of the sulfur atoms from two different Cys residues.

The remaining residues proceeded to side-chain sampling during complex refinement.

### 2.2 Initial Sidechain Conformation Generation

For the first generation, a population of structures was created by sampling rotatable bonds of selected residues. Each dihedral angle *χ* was rotated randomly:

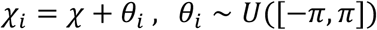

where *χ_i_* represents the dihedral angle of the ith individual in the population, *θ_i_* is a randomly chosen rotation angle within the permissible range, and *U* represents the uniform distribution.

### 2.3 Clash Detection During Sidechain Rotation

Steric clashes were detected during the first generation of sidechain rotations using a detailed clash-detection algorithm. The algorithm examines each atom in the rotated residue against its neighboring atoms. Backbone atoms were excluded from clashing checks if both interacting atoms belonged to the backbone. Hydrogen atoms were also excluded from clash consideration. For each atom in the residue, the algorithm searches for neighboring atoms within a 4.0 Å threshold. A clash was defined when the interatomic distance between any atom pair was less than 1.75 Å, indicating steric overlap. Structures exhibiting at least one such steric clash were flagged and excluded from further consideration, ensuring only feasible conformations proceeded to subsequent docking steps.

Steric clashes between atoms of different residues *C*(*A_i_, A_j_*), were detected using the following criterion:

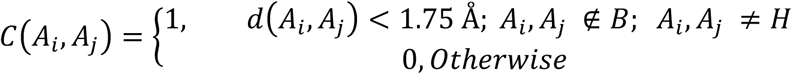

where *d*(*A_i_, A_j_*), is the Euclidean distance between atom coordinates 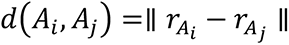 and *B* = {*N,CA,C,O,OXT*}.

A residue *R* was flagged as having a steric clash if at least one clash occurred:

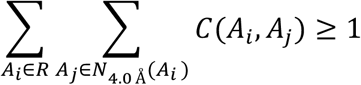

where *N_4.0Å_* (*A_i_*) represents the 4.0 Å neighbor set of the atom *A_i_*. Residues satisfying this criterion underwent further conformational adjustments to resolve the steric clashes which were identified by assessing van der Waals radii and atomic distances in order to preserve steric viability. The stored conformations do not have any steric conflicts. A structure was saved for later processing once all angles had been rotated. Following these rotations, explicit energy minimization was intentionally not carried out because low-energy local minima can be identified through sufficiently dense sampling of sidechain rotamers. Empirically, it was found that a varied ensemble of protein-ligand conformations was produced by randomly or sequentially changing dihedral angles, capturing conformations at local minima.

### 2.4 Docking score evaluation

All rotated structures were then docked using Vina-GPU-2.1 ^17^ and docking scores were computed. Each individual solution *i* in the population be associated with a vector of scores 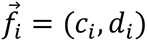 where *c_i_* ∈ ℝ^1^ denotes the clash count for individual *i*, and *d_i_* ∈ ℝ denotes the docking score for individual *i*. To rank individuals by their fitness, a lexicographic order is applied:

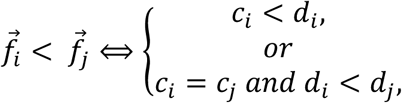

This two-step sorting ensures that clash minimization is the primary criterion, while docking optimization serves as a tie-breaker. After sorting the population *P* = {*i*_1_, *i*_2_, …, *i_n_*} based on the above fitness criteria, a truncation selection is performed:

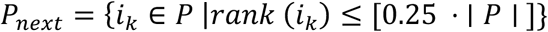

In other words, only the top 25% of individuals, based on the lexicographic ordering of (*c_i_*, *d_i_*), are selected to form the next generation through crossover, mutation, or direct cloning.

### 2.5 GA-Based Refinement of Rotatable Side-Chain Angles

Sidechain conformations were iteratively refined using GA. Selection, crossover, and mutation were all part of the refinement process. The parent pool for the next generation was formed by retaining the top 25% of poses with higher fitness values from each generation, based on their steric feasibility and docking scores. From this pool, two parent structures were selected at random, and the crossover operation was used to inherit the dihedral angles:

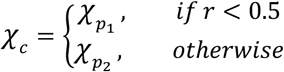

where *χ_c_* is the child’s dihedral angle, 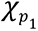 and 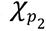 are the corresponding dihedral angles from the two parents, and *r* ∼ *U*([0,1]). Mutations were introduced by perturbing a randomly selected angle:

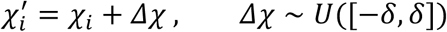

where *δ* is a predefined mutation range that ensures small perturbations in angles without drastic deviations, and *U* is the uniform distribution. We use *δ* = 1 throughout the study. This mutation strategy introduces conformational diversity while preventing premature convergence to suboptimal solutions. Each newly generated conformer was re-evaluated using docking calculations, and the iterative refinement continued for a predefined number of generations. The empirical analysis of docking scores demonstrated that beyond a certain number of sampling steps, the docking score *S_min_*(*N*) converged toward a stable value, confirming the identification of a local minimum.

### 2.6 Cluster-Based Selection of Optimal Protein-Ligand Complex from Iterative Rotational Sampling

The convergence toward the optimal docking pose was enhanced by the recurrent refinement process. Based on steric clash concerns and docking scores, the final best-scoring structure was chosen. By identifying sterically unfavorable interactions and resampling those particular side chains while maintaining the stability of other structural elements, a tailored resampling strategy was put into practice. As a result, the binding conformation might be adjusted without the need for a clear minimization step. After completing all generations, the best complex was selected from the final 10 generations. Poses without steric clashes were filtered, and the top-10 poses per generation based on docking scores were clustered using the root-mean-square deviation (RMSD) of ligand heavy atoms. The cluster with the largest component was identified to ensure structural consistency, and the structure with the best docking score from this cluster was chosen as the final optimized protein-ligand complex. Structures derived via pure rotational sampling were shown to have well-packed sidechains with few steric conflicts and advantageous interactions, frequently requiring no additional modification, according to empirical validation. The technique successfully found near-optimal binding conformations, as evidenced by the best poses showing small changes in geometry and score, when subjected to post hoc energy minimization for validation.

This method maintains computational efficiency while accurately capturing binding poses. By combining iterative refinement with broad conformational sampling, it identifies stable protein–ligand complexes without explicit energy minimization, making it well suited for structure-based modeling and docking optimization. Scheme 1 summarizes the general architecture of the DRGSCROLL.

### 2.7 Hit Rate (Hits%) Calculations

The percentage of true active compounds that are accurately recovered within a specified top-ranked fraction of the screened library is measured by the Hit% (Hit Rate) metric. It gives a clear indication of how well a docking or scoring technique gives active molecules precedence over decoys. All compounds were sorted from most to least favorable according to their projected docking scores in order to determine Hit%. The fraction of real actives in each method’s top N% (e.g., top 1%, 5%, or 10%) of ranked compounds was calculated using the following formula:

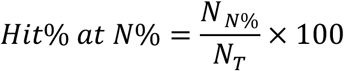

where *N_N%_* is the number of known actives retrieved among the top *N%* of ranked compounds, and *N_T_* is the total number of actives in the dataset.

By putting emphasis on early recognition performance, that is, the number of actives found among the top-ranked candidates most likely to be chosen for experimental validation, this metric enhances enrichment factor (EF) and AUC-based evaluations.

**Scheme 1.**
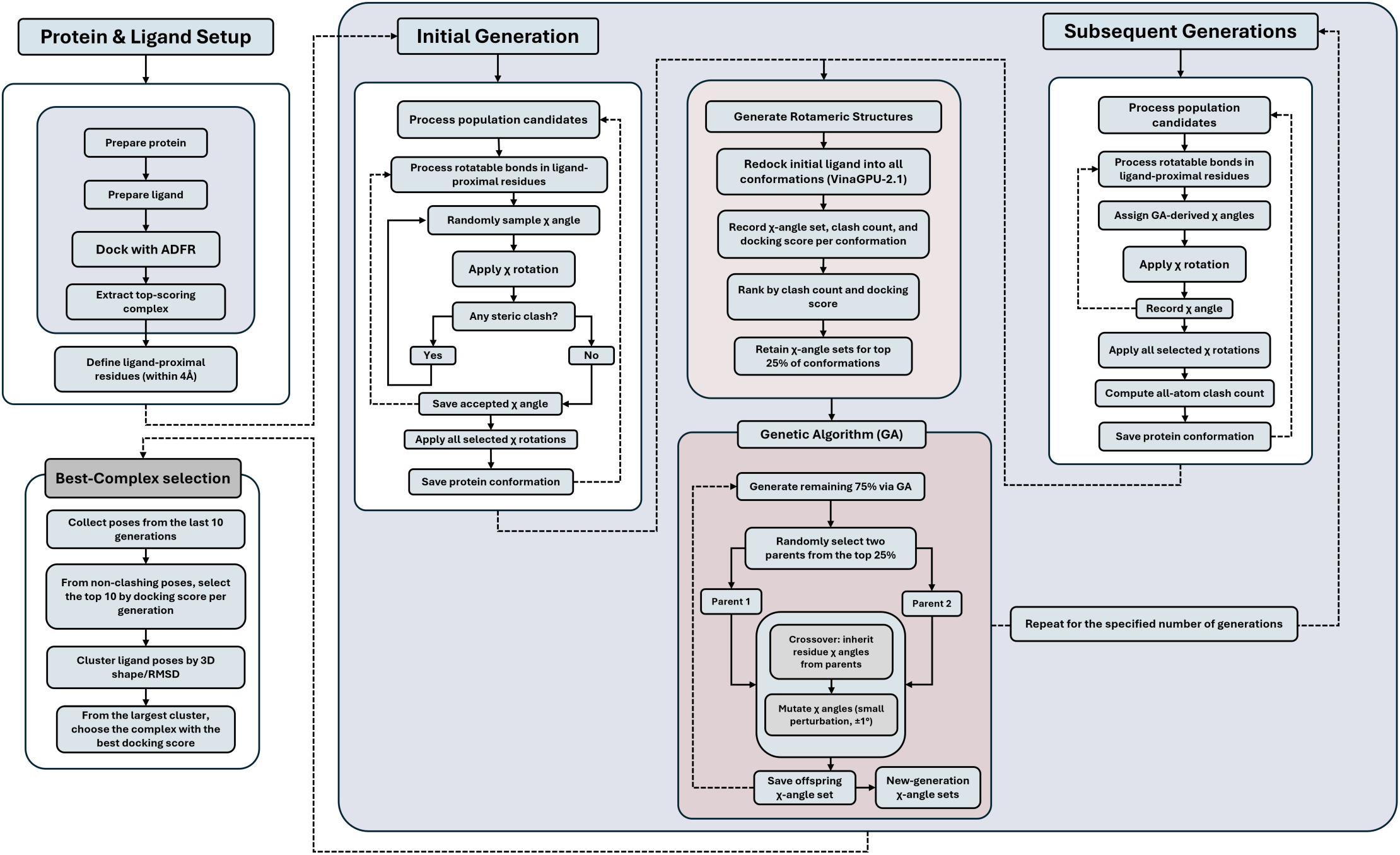
Workflow of the DRGSCROLL side-chain flexible docking algorithm. Protein and ligand are first prepared and docked using ADFR to define binding-site residues. In the first generation, all χ-angles of rotatable side chains around the ligand are systematically sampled, filtered for steric clashes, and docked using Vina GPU 2.1. The top 25% of conformations (based on docking score and clash count) seed the GA stage, where χ-angles are recombined and mutated to explore new side-chain conformations across subsequent generations. The best complex is finally selected by clustering the top structures from the last 10 generations by RMSD and choosing the lowest energy pose from the largest cluster.

## 3. Results and Discussion

### 3.1 Generation-wise convergence of side-chain χ angles and pose quality

In order to rigorously investigate the impact of sidechain flexibility on protein-ligand docking, we utilized DRGSCROLL in this work. For our test set, we chose 50 random co-crystallized protein-ligand complexes from the PDBbind v2018 revised dataset. (https://www.pdbbind-plus.org.cn) Figure S1 shows number of randomly used proteins from each protein family/class. 50 co-crystallized random set forms from 12 different protein classes. A thorough evolutionary search for the best sidechain conformations that improve docking accuracy and stability was made possible using the DRGSCROLL algorithm, which was run over 50 generations with 100 populations per generation (5000 poses in total). Over the course of several generations, the system’s iterative evolutionary algorithm significantly improved docking results by continuously tweaking rotatable sidechains to enhance the binding pocket.

We examine the behavior of the rotatable bonds of the binding pocket residues across the 100 populations per generation in Figure 1A, which focuses on the last, 50^th^ generation. It would indicate that the algorithm converged on particular energetically favorable conformations if the side-chain rotations were limited to a small range. The exploration of side chains of the full 360° range, on the other hand, would suggest that DRGSCROLL permits extensive conformational exploration, which would enable ligands to find ideal positions that would not have been possible otherwise.

**Figure 1.**
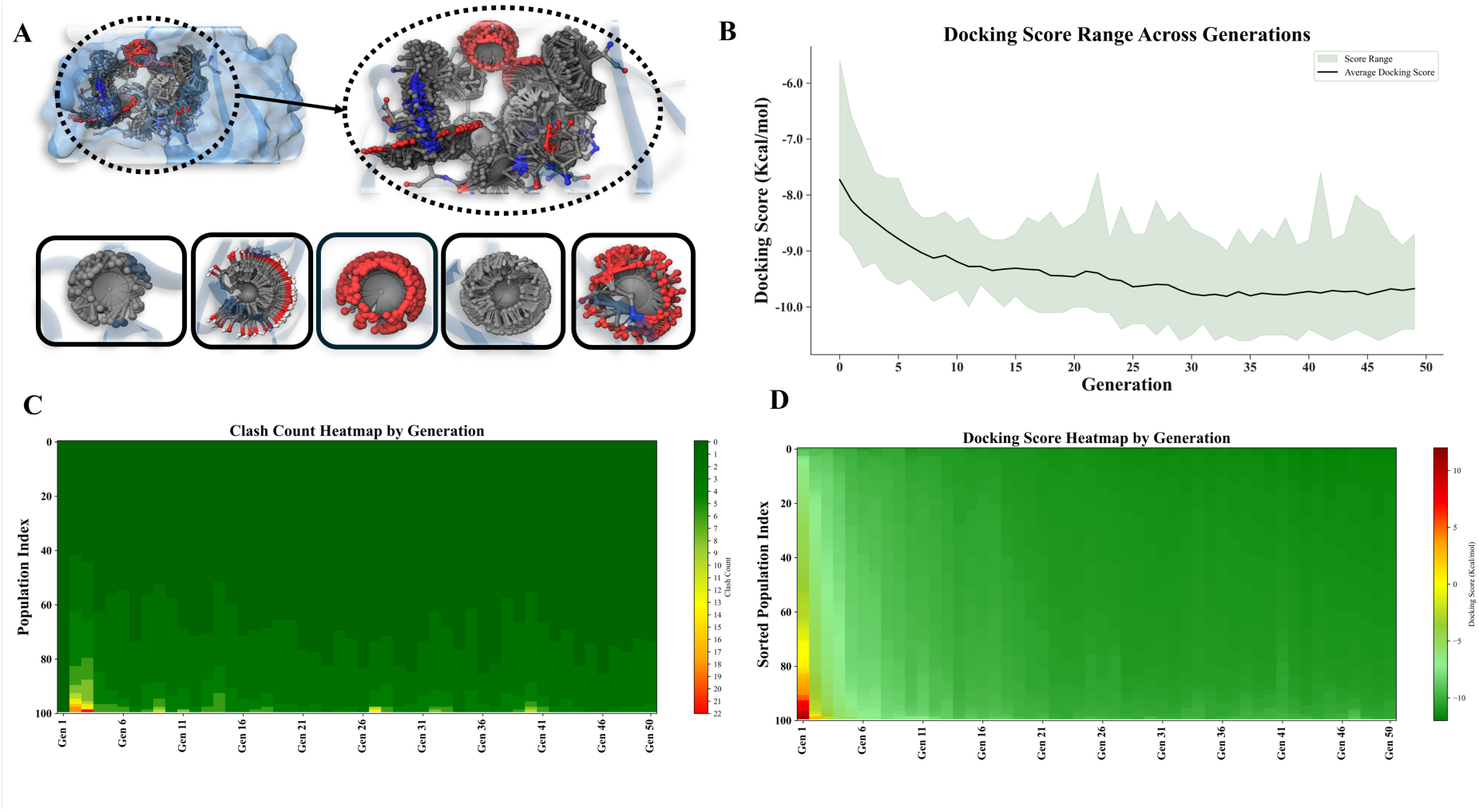
Evolutionary docking optimization with DRGSCROLL. **(A)** Over generations, the binding pocket is improved by iterative sidechain modifications. **(B)** As binding modes improve, average docking scores gradually decline. **(C)** A sharp decline in clash counts during early optimization suggests fewer steric conflicts. **(D)** Docking score heatmaps verify that sidechain flexibility allows for the exploration of a greater conformational space, which results in ligand postures that are more stable and low energy. Panels B–D show representative results, demonstrating docking-score convergence and a declining number of clashes across generations, using the PDBbind structure 1NHZ as the example.

The range of docking scores across generations is shown in Figure 1B, which shows how ligand binding gets better over time as the algorithm advances. The black line indicates the average docking score for each generation, and the shaded area shows the distribution of docking scores. The first generations show a greater variety of docking scores, with some ligands showing comparatively low binding affinities, as the figure illustrates. But as the generations increases, the average score gradually drops and the overall docking scores get better, suggesting improved ligand stability and predicted binding affinity. By improving ligand poses and side-chain flexibility in the binding pocket, the shaded zone narrows, indicating that the algorithm converges towards more advantageous binding poses. This trend demonstrates how well DRGSCROLL refines docking solutions through evolutionary optimization, resulting in complexes that are more stable and energetically advantageous.

The number of complexes devoid of steric clashes steadily rose over generations, as seen in Figure 1C. Initially, because to poor sidechain configurations, the randomly produced conformations frequently led to steric clashes. The selection pressure, however, favored conformations with fewer conflicts as the evolutionary algorithm went on, thereby improving the binding pocket geometry to accommodate ligands more effectively. This pattern demonstrates DRGSCROLL’s capacity to recognize sterically advantageous sidechain conformations, which is essential for obtaining stable and realistic protein-ligand interactions.

The significance of using sidechain flexibility in docking studies, a trait that is sometimes overlooked in conventional docking algorithms, is demonstrated by DRGSCROLL’s capacity to gradually eliminate steric hindrances. The trend in docking scores over the course of evolution is also seen in Figure 1D. As sidechain flexibility was added to the search, the ligands found more energetically advantageous binding modes, as seen by the steady decline of docking scores over subsequent generations. This enhancement implies that stiff binding pockets frequently fall short of offering the best conditions for ligand accommodation. DRGSCROLL allows ligands to explore a larger conformational space by dynamically modifying the sidechains, which ultimately leads to the discovery of more stable and low-energy binding modes. The idea that local sidechain rearrangements can have a substantial effect on ligand location and affinity is further supported by the observed decline in docking scores.

It is interesting to note that although more sidechain flexibility helped many ligand-pocket pairings, some instances only slightly improved docking scores even after several generations. This finding implies that some binding pockets might be intrinsically less compatible with their ligands, even in the face of sidechain flexibility. The inability to obtain appreciable score reductions in these situations suggests that sidechain rotation is not always sufficient to correct for geometric or physicochemical incompatibilities between the ligand and the pocket. This result is consistent with other research that emphasized the significance of pre-existing binding site complementarity rather than making the assumption that flexibility by itself may always improve ligand accommodation. ^18^

The efficiency of DRGSCROLL in improving protein-ligand binding geometries is strongly supported by the growing number of clash-free complexes (Figure 1C) and the improved docking scores over generations (Figure 1D).

This thorough investigation of the rotational space guarantees that the flexibility of DRGSCROLL offers a way to maximize interactions, even in cases when a binding pocket is not initially a good fit for a particular ligand. This leads to better docking scores for certain ligands, but for others, the lack of improvement can be a symptom of a fundamental mismatch with the binding site. The method enables a more comprehensive search for the ideal ligand location by permitting each side chain to spin freely throughout several generations.

### 3.2 DRGSCROLL outperforms conventional engines in interaction fidelity and pose quality

For the same 50 PDB structure set that were taken from the PDBbind database, we also used Vina GPU 2.1, ADFR, Glide, and Induced Fit Docking (IFD) to test and contrast the DRGSCROLL method with alternative docking techniques. In comparison to well-established docking approaches, this research sought to assess the effects of side-chain flexibility introduction in DRGSCROLL on docking performance. The evolutionary optimization of docking solutions exhibited a consistent trend toward lower binding energies across successive generations (Figure S2). As visualized by the viridis heatmap, strong-binding conformations (≤ –5 kcal mol⁻¹) became increasingly dominant after the early generations, reflecting efficient exploration and exploitation of the conformational search space by the GA.

The score distributions among docking methods are depicted in the kernel density estimate (KDE) plots (Figure 2A), which demonstrate that DRGSCROLL samples low-energy binding poses more frequently than conventional techniques. Both IFD and DRGSCROLL obtain lower median docking scores than the other approaches, as seen by the Whisker-box plots (Figure 2B), suggesting that they are better at capturing advantageous binding modes. The advantage of dynamically sampling sidechain flexibility during docking is demonstrated by DRGSCROLL, which displays a larger and denser distribution toward favorable energies, but IFD achieves the lowest median.

**Figure 2.**
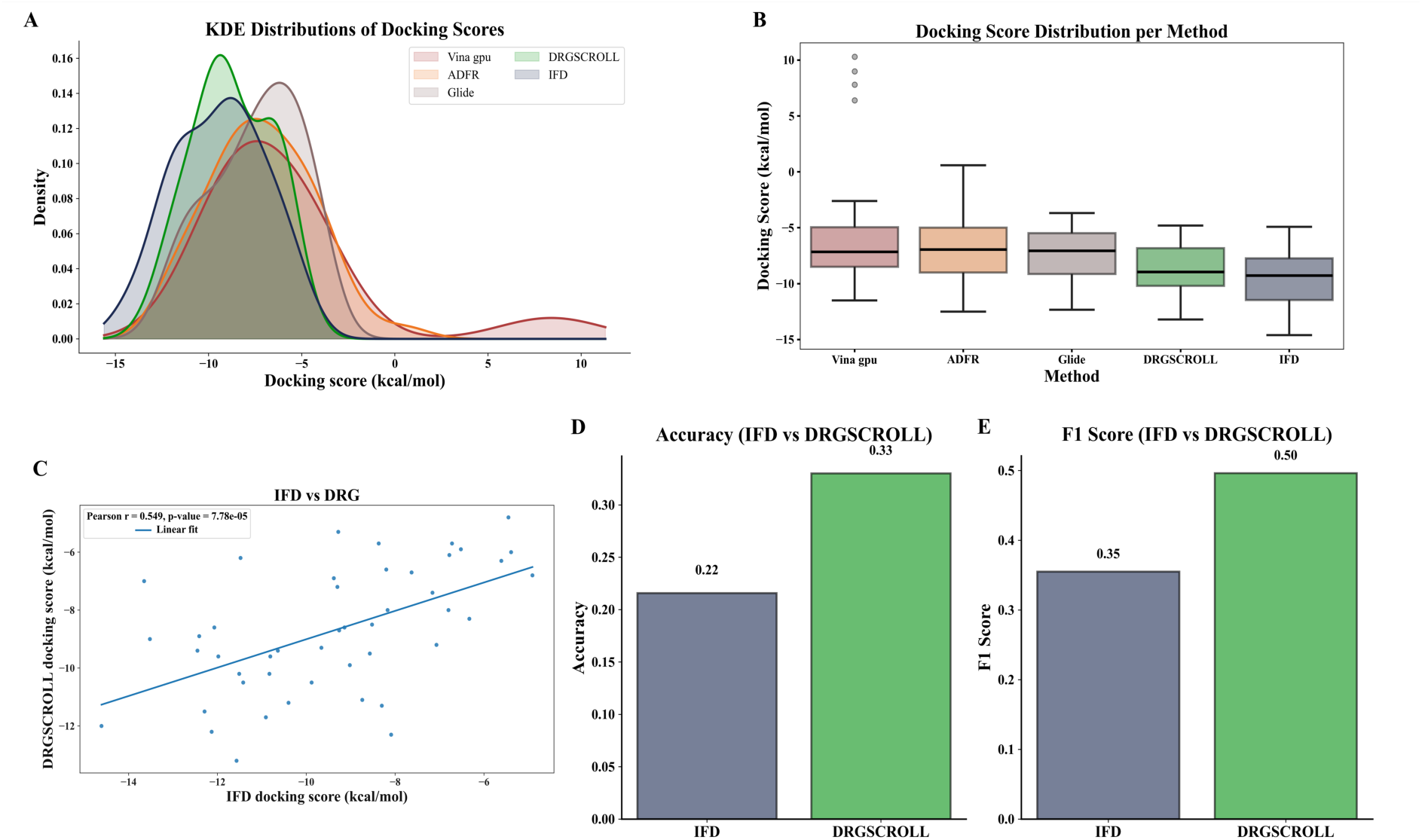
Comparative analysis of DRGSCROLL performance and docking techniques. **(A)** KDE plots display the score distributions for each approach; Vina, ADFR, and Glide are less desirable (lower-energy) poses than DRGSCROLL and IFD sampling. **(B)** Score distributions by approach are shown in box-plots. While DRGSCROLL has a denser spread toward favorable energies, demonstrating the advantage of sidechain flexibility, both IFD and DRGSCROLL attain lower medians than the other methods. **(C)** The correlation between DRGSCROLL and IFD docking scores (Pearson r = 0.589, p < 0.0001) shows that DRGSCROLL can further optimize binding modes while also showing general consistency. **(D, E)** Performance metrics show that DRGSCROLL performs better than IFD in terms of accuracy (0.33 vs. 0.22) and F1 score (0.50 vs. 0.35), indicating that it is more capable of accurately detecting active ligand–protein interactions.

Additionally, compared to alternative approaches, DRGSCROLL exhibits less variability in docking scores, suggesting a more uniform optimization procedure across various ligand-protein complexes. Conventional rigid docking techniques, such Vina, ADFR and Glide, frequently make the assumption that the binding pocket is static, which restricts their capacity to accept ligands that need side-chain orientation changes in order to bind optimally. DRGSCROLL, on the other hand, customizes the binding site environment for every ligand, increasing the possibility of finding advantageous interactions.

IFD and DRGSCROLL’s docking scores show a strong association (Pearson r = 0.589, p < 0.0001), as shown in Figure 2C. In IFD, pocket flexibility is not truly present during the pose search itself. Glide first docks to a largely rigid receptor to select a best initial pose, then, within a user-defined distance around the ligand, local MM optimization of nearby residues and the ligand is performed, and finally the ligand is re-docked into this refined pocket; this yields only partial, localized flexibility and makes outcomes sensitive to the initial Glide pose and the chosen refinement shell. In DRGSCROLL, flexibility is built into the search from the start: A GA co-optimizes ligand placement with on-the-fly side-chain χ-angle rotations across generations, using rotamer-aware, clash-controlled moves. Consequently, DRGSCROLL explores a much larger conformational space, naturally capturing side-chain gating and induced-fit effects without a separate refinement stage, reducing dependence on initial poses. Given that IFD models side-chain flexibility (albeit partially) and its docking scores correlate strongly with DRGSCROLL’s, we used both methods to rigorously assess protein–ligand interaction fidelity. Using PLIP ^19^, we first extracted the experimentally observed residue-level interactions from the reference PDB complex’s co-crystal structure and classified them into four categories: salt bridges, π–π stacking, hydrogen bonds, and hydrophobic interactions. This allowed us to compare the accuracy of each method. For the docked positions that IFD and DRGSCROLL produced, we were able to measure the degree to which each approach replicated the binding network observed in the experiment by mapping each predicted contact onto the reference interaction set according to residue identity and interaction type (Supplementary data).

Recall, interaction precision, and their F1 score were computed for each approach. In addition, the ratio of correctly retained contacts to all experimental interactions was used to determine the overall accuracy of interaction preservation. With a greater accuracy (0.33 vs. 0.22; Figure 2D) and F1 score (0.50 vs. 0.35; Figure 2E), the results showed that DRGSCROLL performed better than the IFD, indicating a stronger agreement with the experimental binding mode. Thus, DRGSCROLL improves docking scores and reduces steric clashes while preserving experimentally observed residue–ligand contacts, indicating that it does more than optimize energetics; it better maintains biologically meaningful, residue-level binding connections. These findings emphasize the advantage of using evolutionary strategies combined with flexible docking to optimize ligand binding. The superior performance of DRGSCROLL in docking score distributions further support the necessity of incorporating adaptive, physics-informed methodologies into modern docking algorithms to achieve more reliable and biologically relevant predictions.

Table S1 shows the docking scores for different protein structures identified by their respective PDB IDs using 5 different docking methods: Vina-GPU 2.1, ADFR, Glide, IFD, and DRGSCROLL. When compared to Vina-GPU 2.1, ADFR, and Glide; DRGSCROLL and IFD generally show consistently lower (more favorable) docking scores across a large number of entries. Notably, several structures, such as 1BXO, 1ERB, 1LEE, and 3SU2, have no data for the IFD approach (shown as *nan*), which suggests that IFD can not find proper docking pose for these co-crystallized structures. The overall variability and distribution of scores emphasize the disparities in the sensitivities and predictive capacities of the docking techniques used, highlighting the significance of choosing the right techniques based on the particular protein-ligand system being studied.

### 3.3 Prospective-style screens on RET TK and PARP1 deliver higher AUROC/AUPRC and early enrichment

Discriminating active hits from inactive compounds is essential for assessing a docking method’s predictive value. We benchmarked DRGSCROLL on two therapeutically relevant targets, RET TK and PARP1, using compound panels curated from ChEMBL with experimentally reported activities. For each target, 100 compounds were docked with (i) Vina-GPU 2.1, which serves as DRGSCROLL’s rigid-receptor baseline engine, and (ii) DRGSCROLL, which augments the same search with side-chain flexibility and GA-based χ-angle optimization. To rigorously test how well the docking methods discriminate actives from inactives, we deliberately curated a bimodal affinity set—overrepresenting very strong binders (pIC₅₀ > 8) and weak binders (pIC₅₀ = 4–6) while intentionally minimizing mid-affinity compounds. Consequently, the pIC₅₀ distribution shows a pronounced split between low- and high-affinity ligands with few intermediates (Figure S3), enriching the dataset at the extremes of potency to sharpen the discrimination assessment.

All docking parameters (grid extents, exhaustiveness, and ligand settings) were held constant across methods; only the treatment of receptor side chains differed. Because docking scores are polarity-consistent, we summarized performance using threshold-free metrics-receiver-operating characteristic (ROC) and precision–recall (PR) curves (Figures 3 and 4) and thresholded metrics (precision, recall, F1, and accuracy; Tables 1 and 2). For the latter, a per-target, per-method decision threshold was chosen to maximize F1 on the docked set without ad-hoc enrichment cutoffs, and uncertainty was quantified via nonparametric bootstrap (1,000 resamples, 95% CIs) with threshold selection repeated within each resample. This design isolates the incremental value of side-chain flexibility and evolutionary optimization relative to the Vina baseline while providing both rank-based (AUROC/AUPRC) and decision-based summaries of predictive performance.

**Figure 3.**
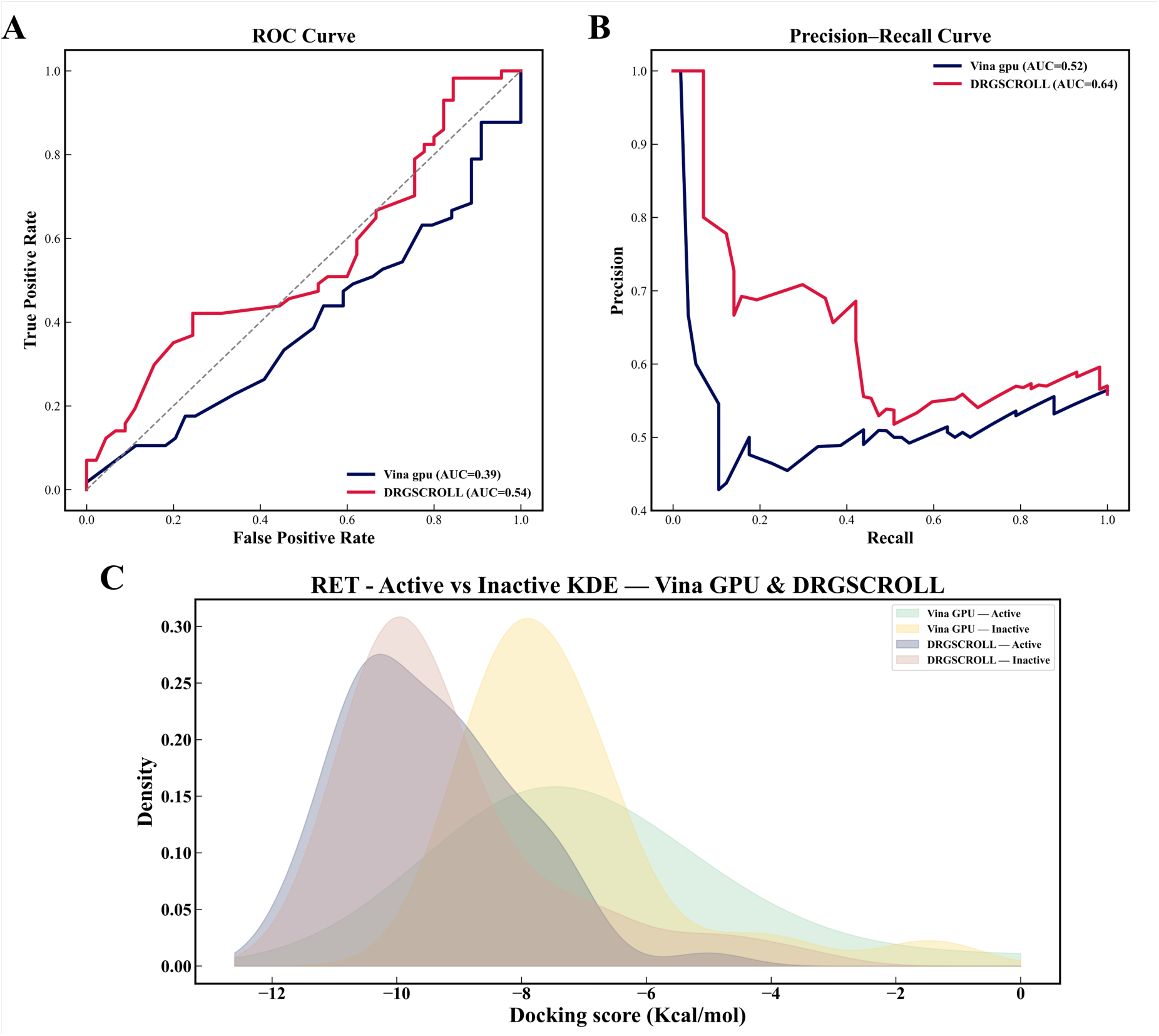
A comparison between Vina GPU 2.1 and DRGSCROLL’s RET docking performance. **(A)** The ROC curve indicates that DRGSCROLL performs better in discriminating between active and inactive ligands, achieving a higher AUC (0.54) than Vina GPU 2.1 (0.39). **(B)** The precision-recall curves verify that DRGSCROLL has a greater precision at comparable recall (AUC = 0.64), compared to Vina GPU 2.1 (AUC = 0.52). **(C)** KDE distributions show that DRGSCROLL can produce more energetically consistent binding modes by better separating active and inactive docking scores.

**Figure 4.**
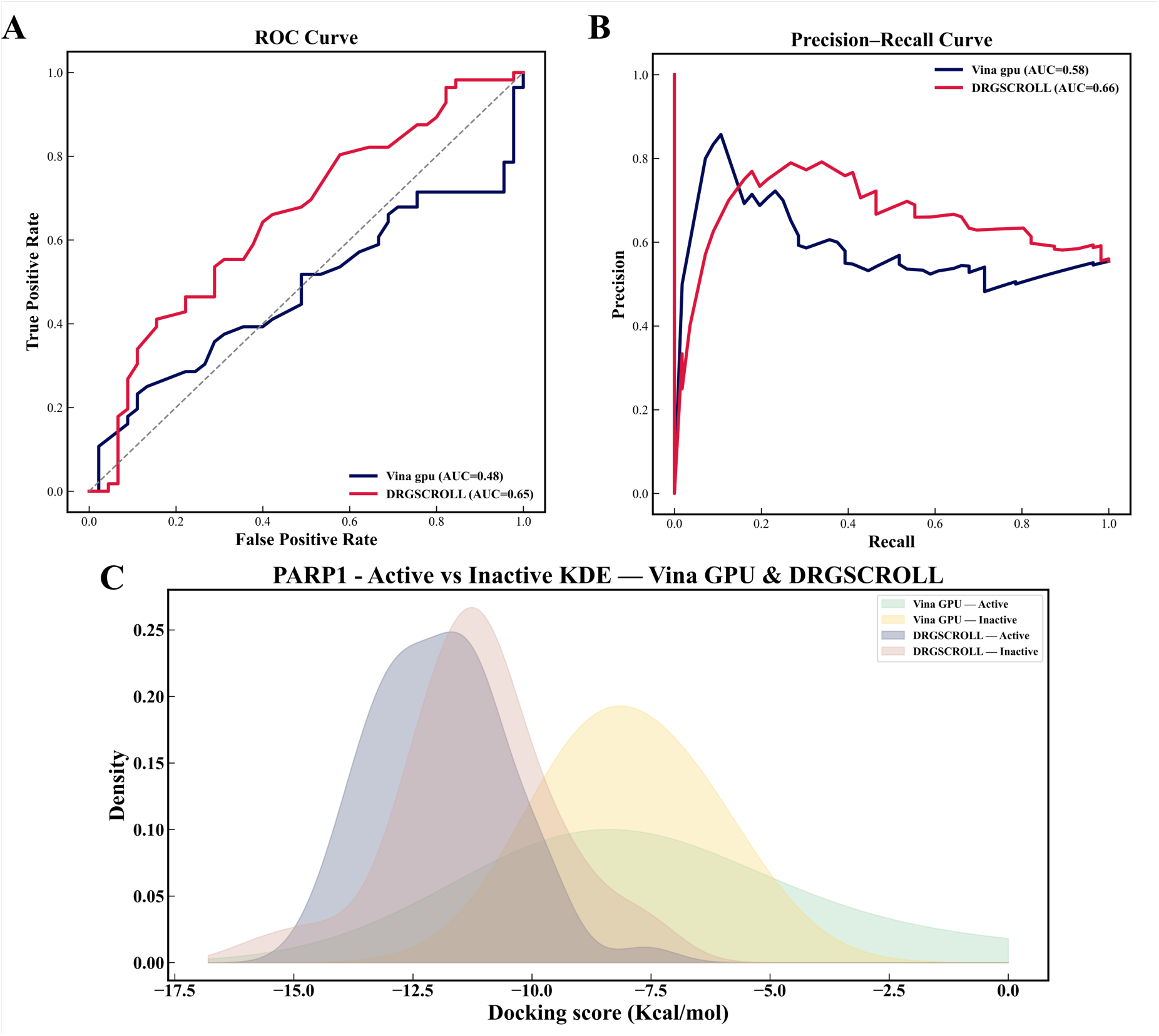
Vina GPU 2.1 and DRGSCROLL’s PARP1 docking performance comparison. **(A)** ROC curve demonstrating DRGSCROLL’s superior detection of active ligands, with a higher AUC (0.65) than Vina GPU 2.1 (0.48). **(B)** According to precision-recall curves, DRGSCROLL outperforms Vina GPU 2.1 in terms of precision-recall balance (AUC = 0.66 against AUC = 0.58). **(C)** KDE charts highlight the advantages of sidechain flexibility by demonstrating how DRGSCROLL more successfully distinguishes between active and inactive docking scores.

**Table 1.** RET docking classification metrics with Vina GPU 2.1 and DRGSCROLL. Once more surpassing Vina GPU 2.1, DRGSCROLL maintains a more balanced trade-off between precision and recall while attaining greater F1 (0.74 vs. 0.72) and accuracy (0.62 vs. 0.56).

|  | Precision | Recall | F1 Score | Accuracy Score |
| --- | --- | --- | --- | --- |
| Vina GPU 2.1 | 0.56 | 1.00 | 0.72 | 0.56 |
| DRGSCROLL | 0.60 | 0.98 | 0.74 | 0.62 |

**Table 2.** Classification metrics for PARP1 docking with DRGSCROLL and Vina GPU 2.1. In comparison to Vina GPU 2.1, DRGSCROLL exhibits better F1 (0.74 vs. 0.71) and accuracy (0.61 vs. 0.55) with balanced precision and recall, while Vina attains recall (1.00) at the expense of more false positives.

|  | Precision | Recall | F1 Score | Accuracy Score |
| --- | --- | --- | --- | --- |
| Vina GPU 2.1 | 0.55 | 1.00 | 0.71 | 0.55 |
| DRGSCROLL | 0.59 | 0.98 | 0.74 | 0.61 |

DRGSCROLL outperforms Vina GPU 2.1, attaining better True Positive Rates (TPR) at lower False Positive Rates (FPR), as seen by the ROC curves (Figures 3A and 4A). AUC values for RET and PARP1 were obtained quantitatively by DRGSCROLL at 0.54 and 0.65, respectively, while Vina GPU 2.1 produced AUC values of 0.39 and 0.48. This indicates that DRGSCROLL provides better predictive reliability, particularly evident for PARP1. Likewise, the PR curves (Figures 3B and 4B) demonstrate that DRGSCROLL reduces false positives while preserving the capacity to identify genuine active chemicals, maintaining greater precision over a wide range of recall levels.

KDEs further highlight the difference between active and inactive molecules (Figures 3C and 4C). By using side-chain flexibility and evolutionary optimization, DRGSCROLL is able to sample more realistic binding modes, as evidenced by the more diverse and well-separated score distributions it generates in comparison to Vina GPU 2.1. DRGSCROLL is significant because it demonstrates that active molecules are moved toward lower (more advantageous) energies, making the distinction between them and inactives more obvious. Vina GPU 2.1, on the other hand, has a limited capacity to distinguish between the two categories, producing wider, more diffuse active distributions with significant overlap with inactives. The classification metrics reinforce these findings. For RET TK (Table 1), DRGSCROLL achieved an F1 score of 0.74 and accuracy of 0.62, compared to 0.72 and 0.56 with Vina GPU 2.1. For PARP1 (Table 2), DRGSCROLL again outperformed Vina GPU 2.1 with an F1 score of 0.74 and accuracy of 0.61, versus 0.71 and 0.55. Although Vina GPU exhibited slightly higher recall (1.00), this came at the cost of lower precision and higher false positives, whereas DRGSCROLL offered a more balanced trade-off between precision and recall.

In sensitivity analyses that excluded method-specific positive (>0) and missing (NaN) docking scores and used pIC_50_ > 6.5 to define activity, we re-evaluated performance with a 1,000-replicate nonparametric bootstrap to derive 95% percentile confidence intervals (CIs). Under this protocol, DRGSCROLL outperformed Vina GPU 2.1 on both RET TK and PARP1, demonstrating higher precision, recall, F1, and accuracy. (Table 3) These results indicate a robust and reproducible performance advantage for DRGSCROLL that persists after score cleaning and uncertainty quantification.

**Table 3.**
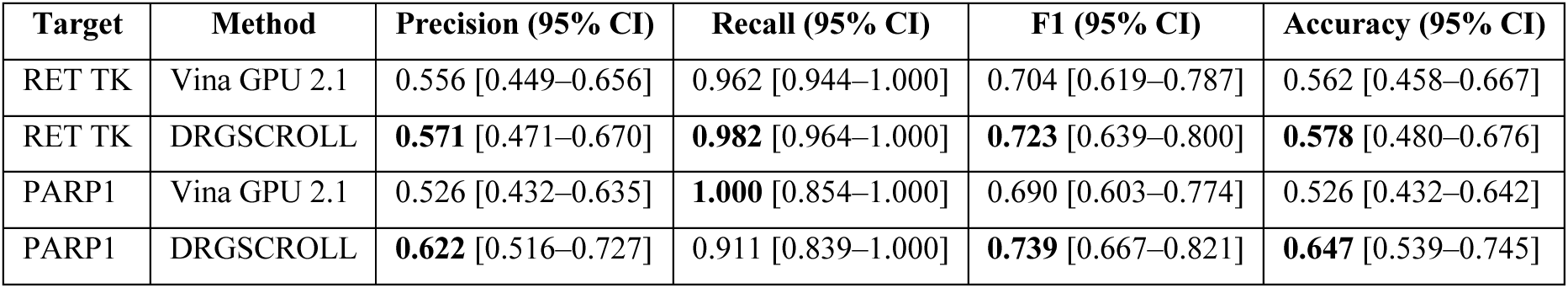
Performance of Vina GPU 2.1 and DRGSCROLL on RET TK and PARP1. Precision, Recall, F1, and Accuracy at the F1-optimal score threshold, with 95% bootstrap CIs (n = 1000). Labels: pIC_50_ > 6.5 = active. Pre-processing: method-specific removal of positive and NaN scores. Score polarity: more negative = better.

Across both targets, DRGSCROLL yielded higher F1 than Vina GPU 2.1 at their respective F1-optimal thresholds (RET: 0.723 vs 0.704; PARP1: 0.739 vs 0.690), with also accuracy scores (RET TK: 0.578 vs 0.562; PARP1: 0.647 vs 0.526). The precision gain is most pronounced on PARP1 (DRGSCROLL 0.622 vs Vina 0.526) where Vina’s threshold achieved perfect recall (1.000) by predicting nearly all compounds as active, increasing false positives and lowering accuracy, while DRGSCROLL traded a modest amount of recall (0.911) for substantially fewer false positives. On RET TK, both methods operated in a high-recall regime (0.96–0.98) with slightly better precision and F1 for DRGSCROLL. Bootstrap 95% CIs generally overlap but are shifted in favor of DRGSCROLL.

When combined, these findings show that DRGSCROLL offers a more precise and physiologically appropriate method of differentiating between ligands that are active and those that are not. DRGSCROLL improves the core Vina GPU 2.1 method by dynamically integrating side-chain flexibility, which results in improved classification performance and more accurate binding predictions for RET TK and PARP1.

Tables S2 and S3 provide the detailed data underlying the ROC curve analyses for RET TK and PARP1 targets, respectively. The ROC curves generated from these data (shown in Figures 3 and 4) evaluate the performance of the docking methods in classifying active and inactive compounds.

### 3.4 Active–decoy benchmarking (DUDE-style): early enrichment and ranking power

In order to choose a representative and chemically diverse subset of active compounds from the 508 actives in the DUDE-PARP1^20^ dataset, we used the RDKit and scikit-learn frameworks to build a clustering-based selection method. In order to obtain binary bit-vectors that represented chemical substructures, all active molecules were first transformed into ECFP4 molecular fingerprints (radius = 4, 4096 bits). These fingerprints were grouped using Euclidean distance and K-Means clustering (k = 5) in the high-dimensional feature space. To guarantee intra-cluster diversity, four representative molecules were chosen from each cluster. A Tanimoto-based Max–Min method, which iteratively selects molecules that maximize their minimum pairwise distance (1 – Tanimoto similarity) from previously chosen members, was used for selection within each cluster. This method prevents redundancy among structurally similar molecules while guaranteeing that the chosen set covers the entire chemical diversity of the active area. 20 different active compounds in all were chosen from the 5 clusters. To verify low redundancy among selected individuals, their structural similarity was assessed by calculating pairwise Tanimoto coefficients and displayed using a heatmap.

The 5 K-Means clusters are shown in the principal component analysis (PCA) projection of ECFP4 fingerprints (Figure S4A), where each hue denotes a different chemical subspace. The 20 chosen active molecules are shown by yellow star markers, and their uniform distribution throughout clusters attests to the efficacy of diversity sampling. Further evidence that the selected actives are structurally non-redundant and collectively capture the general structure of the DUDE-PARP1 active chemical space is provided by the pairwise Tanimoto similarity heatmap (Figure S4B), which also shows low intra-selection similarity (usually Tc < 0.4).

The accompanying Python script we built a property-matched and similarity-restricted technique to select decoy compounds for each of the 20 typical actives found in the DUDE-PARP1 dataset. The first step in the process is creating a candidate decoy pool that corresponds to each active’s physicochemical profile. The Tanimoto similarity between each active and its property-matched decoy candidates is then calculated once all remaining candidates have been represented as extended-connectivity fingerprints (ECFP4; radius 4, 4096 bits). 3 rounds of decoy selection are made based on diminishing similarity windows: a primary range of 0.35 to 0.55, a relaxed range of 0.30 to 0.60 (not including decoys that have already been allocated), and a final cap of Tc < 0.65. A greedy global assignment process makes sure that each active receives up to 10 distinct decoys while avoiding the reuse of the same decoy for numerous actives. Candidate pairs are sorted by similarity inside each run. A balanced library of 200 decoys that are structurally different from their parent actives but have the same properties is produced by this process.

Tables S4 provides detailed data underlying the ROC curve analyses for active versus decoy analysis. The ROC curves generated from these data (shown in Figure 5) evaluate the performance of the docking methods in classifying active and decoy compounds. Pairwise similarity and molecular-space visualization were used to assess the final dataset. The ECFP4 Tanimoto coefficient heatmap displays the similarity distribution of each active’s 10 assigned decoys in order of least to most similar. While staying within the intended similarity restrictions, the majority of values fall between 0.10 and 0.18 (median 0.13), indicating that decoys and actives are chemically different. (Figure S5) A two-dimensional embedding of molecular fingerprints is provided by the accompanying UMAP projections, in which each active is represented by a yellow star around by its ten blue decoys. The decoys always group together in adjacent but non-overlapping areas, suggesting that property similarity is maintained without the need for scaffold duplication. These evaluations collectively show that the selection process effectively produces decoy sets that are structurally unique, property-matched, and chemically viable, making them appropriate for reliable benchmarking of docking and scoring techniques. (Figure S6) For the classification of actives and decoys, the trade-off between true positive rate (sensitivity) and false positive rate is depicted by the ROC curve. (Figure 5A)

**Figure 5.**
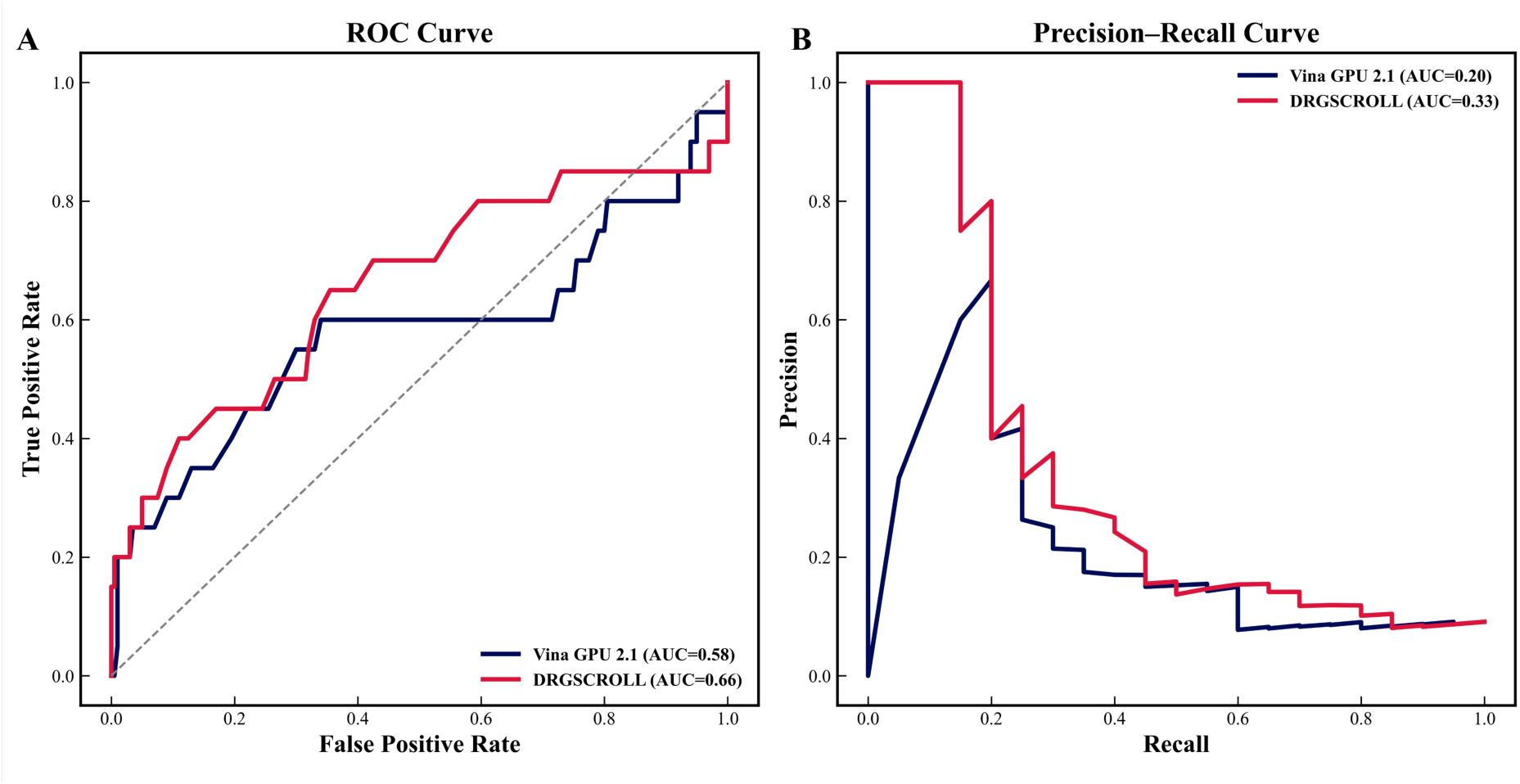
Performance comparison of Vina GPU 2.1 and DRGSCROLL in categorization. **(A)** A ROC curve comparing the true and false positive rates. Precision as a function of recall is displayed in the precision–recall curve **(B)**. In both AUC and early identification measures, DRGSCROLL (red) continuously beats Vina GPU 2.1 (navy), demonstrating its enhanced capacity to differentiate active ligands from decoys.

The AUC value of 0.66 for DRGSCROLL was higher than 0.58 for Vina GPU 2.1, suggesting better overall classification performance and more precise active compound ranking. The PR curve (Figure 5B) also shows how DRGSCROLL has improved in identifying real actives early on. Vina GPU 2.1’s area under the PR curve (AUC-PR) was 0.20, whereas DRGSCROLL’s was 0.33. This indicates that DRGSCROLL maintains a greater level of precision as recall levels increase. Classification metrics have been compared in Table 4. Together, our findings show that DRGSCROLL offers more dependable active prioritization, especially in the top-ranked screening subgroup, corroborating its improved score consistency when compared to traditional docking methods.

**Table 4.** Classification metrics for PARP1-DUDE compounds docking with DRGSCROLL and Vina GPU 2.1.

|  | Precision | Recall | F1 Score | Accuracy | AUC |
| --- | --- | --- | --- | --- | --- |
| Vina GPU 2.1 | 0.14 | 0.55 | 0.22 | **0.65** | **0.58** |
| DRGSCROLL | **0.15** | **0.65** | **0.25** | 0.64 | **0.66** |

The Hit% (True Active Recovery) was calculated across several screening levels (Top 10%, 15%, 20%, 50%, and 75%) in order to assess the docking algorithms’ early identification capabilities. The percentage of known active molecules that are accurately rated in the top section of the screened library is measured by the metric.

Across all datasets, DRGSCROLL consistently achieved higher hit recovery rates than Vina GPU 2.1, indicating stronger prioritization of active compounds. In comparison to Vina GPU 2.1, which had Hit% values of 12.3%, 19.3%, and 24.6% at the Top 10%, 15%, and 20% levels, DRGSCROLL obtained Hit% values of 14.0%, 19.3%, and 26.3%, respectively. (Figure 6A) The early retrieval zone (Top 10%) showed the greatest relative improvement in DRGSCROLL, demonstrating its enhanced ability to discriminate between strong binders.

**Figure 6.**
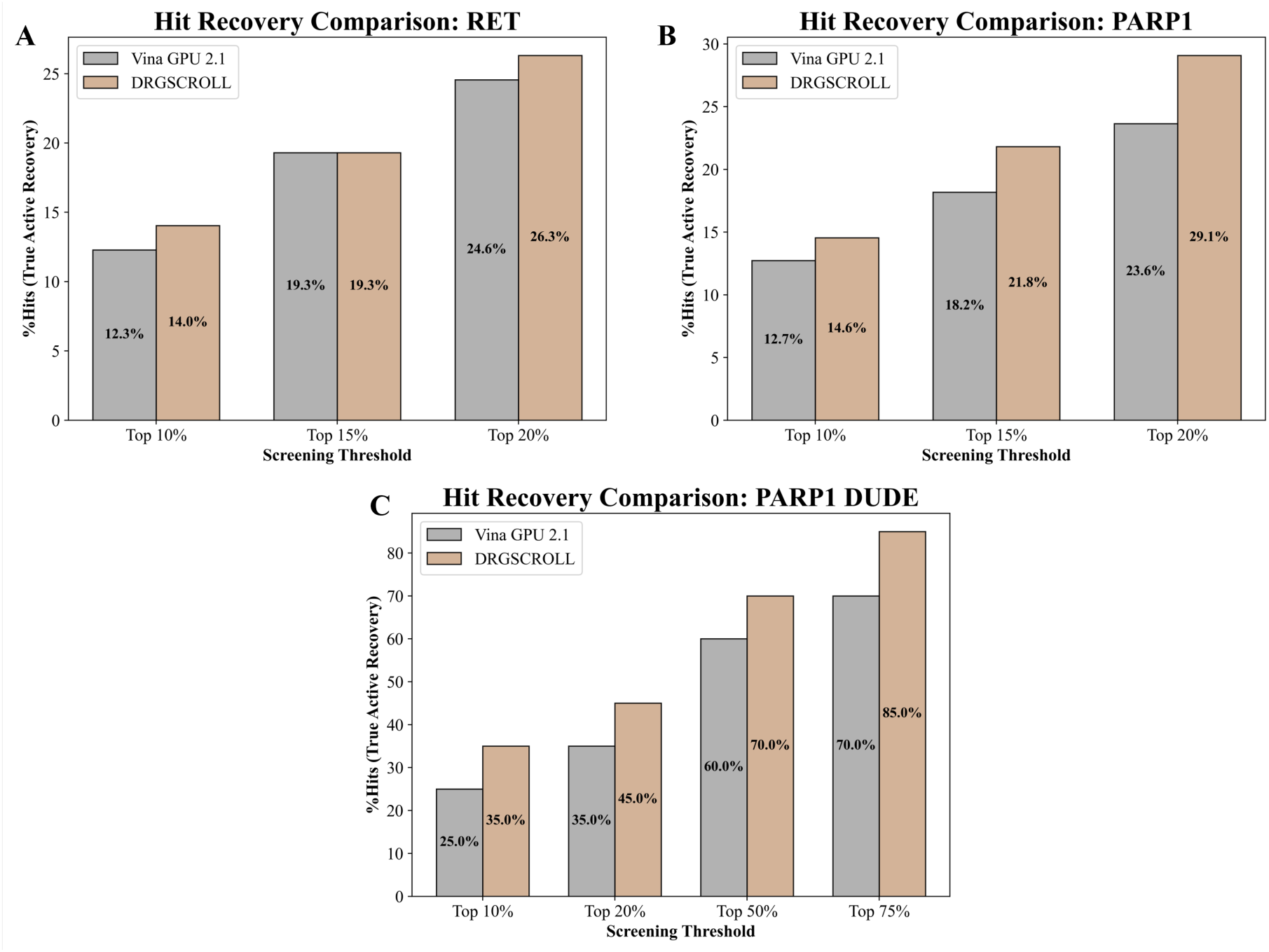
A comparison of Vina GPU 2.1 and DRGSCROLL’s hit recovery. RET, PARP1, and PARP1 DUDE datasets (A, B, and C, respectively). The percentage of known actives that were successfully recovered inside the top N% of the library screened (Hit%) is shown by bars. Higher hit recovery rates across thresholds are routinely attained using DRGSCROLL, suggesting better active compound prioritization and early enrichment.

DRGSCROLL recovered 14.6%, 21.8%, and 29.1% of actives compared to 12.7%, 18.2%, and 23.6%, respectively, surpassing Vina GPU 2.1 once more across all thresholds. (Figure 6B) According to these findings, early hit retrieval increased by about 24%, indicating improved scoring sensitivity for PARP1 complex actives.

The pattern was validated on a bigger scale with 20 actives and 200 decoys per active. At the Top 10%, 20%, 50%, and 75% thresholds, DRGSCROLL obtained Hit% values of 35.0%, 45.0%, 70.0%, and 85.0%, respectively, while Vina GPU 2.1 achieved 25.0%, 35.0%, 60.0%, and 70.0%. (Figure 6C) In the early discovery phase (top 10% and 20%), the improvement was especially noticeable, confirming DRGSCROLL’s outstanding early-enrichment capacity.

Overall, DRGSCROLL consistently produces stronger true-active recovery across all thresholds, as seen by the Hit% comparison across the RET, PARP1, and DUDE ^20^ datasets, demonstrating its improved ranking power and scoring robustness.

## 4. Conclusions

Structure-based docking remains constrained by how receptors are modeled. DRGSCROLL addresses a long-standing gap by coupling ligand pose search with continuous optimization of side-chain χ angles inside a GA loop, balancing interaction energy with explicit clash penalties and intentionally omitting per-iteration minimization to prioritize exploration. Across 50 PDBbind complexes run for 50 generations × 100 populations, this design produced monotonic reductions in clash counts and progressive improvements in docking energies, indicating convergence to sterically viable, low-energy pocket conformations that rigid-receptor protocols seldom access. Against widely used engines (Vina, ADFR, Glide) and IFD; DRGSCROLL sampled more favorable energy distributions with lower medians on the same set of protein-ligand complexes, mirroring the benefits of induced fit while avoiding heavy local minimization. Kernel-density and box-plot comparisons show DRGSCROLL’s distributions shifted and tightened toward lower energies, and its scores correlate with IFD, yet DRGSCROLL better preserved residue-level contacts from co-crystal references. These observations support the central premise that modeling side-chain plasticity during the search, rather than as a post-hoc tweak, improves both physical plausibility and binding-mode fidelity.

In prospective virtual-screening style tests on RET TK and PARP1, using ChEMBL-derived actives/inactives and DUDE property-matched decoys, DRGSCROLL improved early-retrieval behavior over its underlying rigid baseline (Vina-GPU-2.1). Classification metrics likewise favored DRGSCROLL, reflecting a more balanced precision–recall trade-off than the high-recall-but-noisy baseline. Collectively, these gains indicate that on-the-fly side-chain adaptation helps separate actives from inactives and yields more realistic binding modes without resorting to expensive receptor relaxations.

### 4.1 Implementation and Accessibility

The DRGSCROLL docking platform and webserver was optimized for GPU-accelerated execution, enabling receptor-flexible docking to be performed at practical timescales. On an NVIDIA RTX 4070 GPU, complexes containing approximately 300–500 residues typically complete around 2-3 min/generation, including the generation and evaluation of side-chain rotamers. Larger systems or those with extensive pocket plasticity may require proportionally longer runtimes, but the algorithm remains scalable across multiple receptor targets. This balance between flexibility and computational efficiency makes DRGSCROLL suitable for both benchmarking studies and medium-throughput virtual screening campaigns.

### 4.2 Limitations and future perspectives

DRGSCROLL currently (i) restricts flexibility to side chains within a 4 Å shell, (ii) keeps the backbone fixed, (iii) relies on Vina-GPU-2.1 scoring during the GA, and (iv) enforces hard clash filters (e.g., 1.75 Å interatomic cutoff; backbone/hydrogen handling) that may exclude rare but resolvable strain states. The moderate AUC (0.54) value of RET TK also highlights that flexibility alone does not fully overcome ligand–site incompatibility; incorporating water networks, alternative protonation/tautomer states, metal coordination, and long-range conformational changes may be necessary for some targets.

Several extensions are natural: (1) selective backbone mobility via normal-mode or elastic-network subspaces; (2) multi-objective GA that co-optimizes hydrogen-bond geometry, desolvation, and interaction fingerprints alongside energy and clashes; (3) consensus/ML rescoring (e.g., GNINA-style CNNs) on GA-generated ensembles combined with physics-informed learning models; and (4) explicit water/protonation sampling during the search. Furthermore, DRGSCROLL may demand long runtimes when screening very large ligand libraries. This is a consequence of its thorough search and evaluation strategy, which prioritizes accuracy over speed. In future versions, we will study and implement optimizations to decrease the running time without compromising ranking fidelity, enrichment, or overall hit-identification success.

Taken together, our results show that embedding continuous side-chain flexibility into the search itself enables DRGSCROLL to recover biologically faithful poses while preserving discovery-scale throughput. Because the method is transparent and readily parameterized, it generalizes to side-chain–gated or shallow pockets and is well suited as a default engine for prospective screening where induced fit is expected. We therefore anticipate that deploying DRGSCROLL upstream of standard refinement workflows will shorten iteration cycles and raise hit quality in real-world discovery campaigns.

## ASSOCIATED CONTENT

The DRGSCROLL docking platform and web server are accessible at https://www.drgscroll.com and https://drgscroll.bau.edu.tr/, respectively. Benchmarking and virtual screening datasets were curated from the PDBbind v2018 database (https://www.pdbbind-plus.org.cn), the ChEMBL database (https://www.ebi.ac.uk/chembl/), and the Directory of Useful Decoys (DUD-E) (https://dude.docking.org/targets/parp1). All data used for the validation of the algorithms have been deposited in Zenodo and are available at https://zenodo.org/records/17733136.

## Supporting Information

The Supporting Information is available free of charge at [].

Benchmark docking scores comparing Vina GPU 2.1, ADFR, Glide, DRGSCROLL, and IFD; detailed docking scores and ROC curve data for RET tyrosine kinase and PARP1; docking scores for PARP1 active versus decoy ligands selected from DUDE, were provided as Supplementary Tables. Distribution of protein families used in the study; heatmaps showing docking score evolution across generations; compound distribution according to binding affinity types; active compound diversity evaluation and cluster-based selection; ECFP4 Tanimoto similarity heatmaps; and UMAP projections of active stars and decoys were also provided.

## AUTHOR INFORMATION

### Corresponding Author

Serdar Durdağı – Lab for Innovative Drugs (Lab4IND), Computational Drug Design Center (HİTMER), Bahçeşehir University, İstanbul, Türkiye; Department of Pharmaceutical Chemistry, Bahçeşehir University, School of Pharmacy, İstanbul, Türkiye; Quantitative System Biology Lab, Faculty of Medicine, Biruni University, Istanbul, 34015, Türkiye; Computational Biology and Molecular Simulations Lab, Department of Biophysics, School of Medicine, İstanbul, Türkiye.

### Authors

Ehsan Sayyah – Lab for Innovative Drugs (Lab4IND), Computational Drug Design Center (HİTMER), Bahçeşehir University, İstanbul, Türkiye; Neuroscience Program, Graduate School, Bahçeşehir University, İstanbul, Türkiye.

Muhammet Eren Ulug – Lab for Innovative Drugs (Lab4IND), Computational Drug Design Center (HİTMER), Bahçeşehir University, İstanbul, Türkiye.

Huseyin Tunc – Department of Biostatistics and Medical Informatics, School of Medicine, Bahçeşehir University, İstanbul, Türkiye.

### Notes

The authors declare no competing financial interest.

## ACKNOWLEDGMENTS

The numerical calculations reported in this paper were partially performed at TÜBİTAK ULAKBIM, High Performance and Grid Computing Center (TRUBA resources). This study was partially funded by the T.C. Istanbul Development Agency, project no. TR10/21/YEP/0133. This study was also funded by the Scientific Research Projects Commission of Bahçeşehir University. Project numbers: BAP.2024.01.42 and BAP.2022.01-12.

## Notes

### Competing Interest Statement

The authors have declared no competing interest.

