## Supporting Information for "DRGSCROLL: Achieving Full Side-Chain Flexibility in Docking Simulations through Genetic Algorithm Framework"

[1]Lab for Innovative Drugs (Lab4IND), Computational Drug Design Center (HİTMER), Bahçeşehir University, İstanbul, Türkiye; [2]Neuroscience Program, Graduate School, Bahçeşehir University, İstanbul, Türkiye; [3]Department of Biostatistics and Medical Informatics, School of Medicine, Bahçeşehir University, İstanbul, Türkiye; [4]Department of Pharmaceutical Chemistry, Bahçeşehir University, School of Pharmacy, İstanbul, Türkiye; [5]Quantitative System Biology Lab, Faculty of Medicine, Biruni University, Istanbul, 34015, Türkiye; [6]Computational Biology and Molecular Simulations Lab, Department of Biophysics, School of Medicine, İstanbul, Türkiye

* (SD)

**Table S1.** Benchmark of docking scores of protein-ligand complexes using five different docking techniques. Vina GPU 2.1, ADFR, Glide SP, DRGSCROLL, and IFD binding scores (kcal/mol) for PDB complexes are compared.

| PDB ID | **VinaGPU_2.1** | **ADFR** | **Glide SP** | **DRGSCROLL** | **IFD** |
|---|---|---|---|---|---|
| 1B6H | -8.5 | -4.2 | -10.8 | -8.9 | -12.4 |
| 1BGQ | -8.4 | -8.4 | -6.4 | -12.3 | -8.1 |
| 1BXO | -9.3 | -6.2 | -5.9 | -10.3 | nan |
| 1C83 | -8.5 | -9.3 | -9.4 | -10.5 | -11.4 |
| 1ERB | -7.8 | -9.0 | -8.9 | -10.1 | nan |
| 1FKG | -8.5 | -6.6 | -7.8 | -10.5 | -9.9 |
| 1HMT | -5.6 | -4.6 | -5.3 | -6.8 | -4.9 |
| 1LEE | -9.7 | -4.9 | -8.1 | -9.2 | nan |
| 1NHZ | -3.3 | -12.5 | -12.1 | -12.0 | -14.6 |
| 2IHJ | -8.1 | -3.2 | -10.4 | -9.6 | -12.0 |
| 3FUZ | -6.1 | -7.4 | -7.3 | -7.2 | -9.3 |
| 3FVK | -9.5 | -9.8 | -7.8 | -9.4 | -10.6 |
| 3G30 | -6.8 | -4.7 | -4.3 | -8.0 | -6.8 |
| 3GBA | -10.3 | -11.2 | -8.7 | -9.6 | -10.8 |
| 3UPV | -3.7 | -1.8 | -6.6 | -5.3 | -9.3 |
| 4ABE | -5.9 | -6.9 | -7.7 | -8.0 | -8.2 |
| 4MMP | -8.4 | -5.3 | -8.1 | -9.3 | -9.7 |
| 4PP3 | -11.3 | -11.4 | -10.4 | -13.2 | -11.6 |
| 4WKN | -8.6 | -9.7 | -11.4 | -9.4 | -12.5 |
| 4XOE | -7.3 | -5.9 | -5.7 | -8.3 | -6.3 |
| 5UOO | -9.4 | -8.9 | -6.3 | -11.7 | -10.9 |
| 6AYI | -9.7 | -9.0 | -10.1 | -10.2 | -10.8 |
| 6CPA | -8.5 | -3.4 | -10.0 | -10.2 | -11.5 |
| 7STD | -10.0 | -8.3 | -8.1 | -11.3 | -8.3 |
| 4AG8 | 10.3 | -11.90 | -11.0 | -8.6 | -12.1 |
| 4AGC | 7.8 | -10.80 | -11.5 | -7.0 | -13.7 |
| 4AGL | -4.6 | -7.00 | -4.6 | -6.0 | -5.4 |
| 4AGM | -4.9 | -8.20 | -4.2 | -6.1 | -6.8 |
| 4AGN | -5.1 | -8.40 | -6.2 | -6.7 | -7.6 |
| 4AGO | -4.3 | -7.90 | -3.7 | -5.7 | -6.7 |
| 4AGP | -5.4 | -8.00 | -5.2 | -6.6 | -8.2 |
| 4AGQ | -5.2 | -9.50 | -5.8 | -6.9 | -9.4 |
| 4AHR | -3.9 | -3.00 | -4.2 | -5.9 | -6.5 |
| 4AHS | -3.7 | -4.00 | -4.0 | -4.8 | -5.5 |

| | | | | | |
|---|---|---|---|---|---|
| 4AHU | -4.4 | -4.30 | -5.1 | -6.3 | -5.6 |
| 4AI5 | -5.6 | -4.50 | -5.1 | -7.4 | -7.2 |
| 4AOI | -11.5 | -10.80 | -11.4 | -12.2 | -12.1 |
| 4DFG | -5.1 | 0.60 | -4.3 | -5.7 | -8.4 |
| 4KNN | -8.0 | -5.80 | -7.2 | -8.6 | -9.1 |
| 4M2R | -6.4 | -6.60 | -5.4 | -8.5 | -8.5 |
| 4MPN | -8.7 | -6.80 | -6.1 | -9.9 | -9.0 |
| 4ZYF | 9.0 | -8.50 | -9.1 | -6.2 | -11.5 |
| 10GS | -7.0 | -4.90 | -6.3 | -8.7 | -9.3 |
| 186L | -7.3 | -5.70 | -7.0 | -9.2 | -7.1 |
| 2R9X | -7.5 | -6.50 | -6.9 | -9.5 | -8.6 |
| 3IOD | -6.1 | -6.90 | -9.2 | -11.5 | -12.3 |
| 3SU2 | -2.6 | -9.90 | -6.6 | -6.4 | nan |
| 4CWF | -9.2 | -8.00 | -7.9 | -11.2 | -10.4 |
| 6EQU | -8.0 | -6.80 | -5.2 | -11.1 | -8.7 |
| 4ASD | 6.4 | -11.6 | -12.3 | -9.0 | -13.5 |

**Table S2.** Docking scores of specific RET tyrosine kinase ligands using the ROC curve. Vina GPU 2.1 and DRGSCROLL were used to dock representative active chemicals with experimentally published pIC50 values.

| Compound | $pIC_{50}$ | Vina GPU 2.1 (kcal/mol) | DRGSCROLL (kcal/mol) |
|---|---|---|---|
| CHEMBL1171837 | 9.52 | 5.3 | -6.00 |
| CHEMBL3355009 | 9.4 | -6.2 | -6.50 |
| CHEMBL3355010 | 9.40 | -5.8 | -6.00 |
| CHEMBL3775169 | 9.39 | -7.2 | -4.60 |
| CHEMBL388978 | 9.34 | -6.5 | -6.30 |
| CHEMBL4062877 | 9.7 | -8.0 | -6.60 |
| CHEMBL4067871 | 10.40 | -7.7 | -6.40 |
| CHEMBL4075917 | 9.4 | -7.8 | -6.10 |
| CHEMBL4105329 | 9.15 | -7.4 | -7.00 |
| CHEMBL4203429 | 9.22 | -4.4 | -4.90 |
| CHEMBL4205243 | 9.4 | -4.8 | -4.10 |
| CHEMBL4212513 | 9.40 | -6.7 | -6.60 |
| CHEMBL4214146 | 9.15 | -5.8 | -5.60 |
| CHEMBL4218127 | 9.4 | -5.7 | -5.70 |
| CHEMBL440727 | 9.12 | -7.4 | -5.30 |
| CHEMBL4582651 | 9.40 | -7.1 | -7.00 |
| CHEMBL4853155 | 9.14 | -8.5 | -5.90 |
| CHEMBL4854822 | 9.27 | -8.1 | -6.40 |
| CHEMBL4870957 | 9.17 | -10.1 | -7.70 |
| CHEMBL1688861 | 9.1 | 3.1 | -6.80 |
| CHEMBL4213626 | 9.1 | -4.6 | -4.9 |
| CHEMBL4846521 | 9.21 | 11.1 | -7.7 |
| CHEMBL4877264 | 9.07 | -8.4 | -6.5 |

| | | | |
|---|---|---|---|
| CHEMBL4580325 | 9 | -4.5 | -4.8 |
| CHEMBL3774904 | 8.96 | -6.9 | -6.1 |
| CHEMBL4791998 | 8.92 | -9.3 | -8.0 |
| CHEMBL1946170 | 8.82 | -1.3 | -5.9 |
| CHEMBL4454732 | 8.77 | -8.0 | -6.1 |
| CHEMBL3775336 | 8.7 | -7.3 | -5.8 |
| CHEMBL1980995 | 8.66 | -5.7 | -6.9 |
| CHEMBL3774489 | 8.59 | -7.0 | -5.2 |
| CHEMBL4846921 | 8.54 | -8.4 | -7.6 |
| CHEMBL3809489 | 8.52 | 4.6 | -9.6 |
| CHEMBL3775879 | 8.49 | -6.9 | -5.5 |
| CHEMBL3671311 | 8.45 | -8.4 | -5.9 |
| CHEMBL3687223 | 8.44 | -7.7 | -4.6 |
| CHEMBL3774580 | 8.41 | -6.9 | -5.8 |
| CHEMBL4116008 | 8.4 | 1.3 | -5.5 |
| CHEMBL3687222 | 8.96 | -8.2 | -6.4 |
| CHEMBL24828 | 9.89 | -4.3 | -5.8 |
| CHEMBL4872605 | 9.44 | -1.3 | -7.7 |
| CHEMBL4559134 | 9.4 | -5.3 | -7.6 |
| CHEMBL4461418 | 8.96 | -8.2 | -5.4 |
| CHEMBL2180602 | 8.89 | -6.9 | -7.1 |
| CHEMBL4859281 | 8.86 | -7.9 | -7.6 |
| CHEMBL4211949 | 7.69 | -6.6 | -6.7 |
| CHEMBL3925190 | 7.67 | -7.8 | -6.7 |
| CHEMBL5075257 | 7.66 | -8.8 | -6.0 |
| CHEMBL3775376 | 7.64 | -6.9 | -7.3 |
| CHEMBL2403370 | 7.6 | -8.1 | -6.4 |
| CHEMBL3913553 | 7.58 | -8.0 | -6.7 |
| CHEMBL3775086 | 7.54 | -8.8 | -7.4 |
| CHEMBL1807521 | 7.42 | -8.0 | -6.0 |
| CHEMBL3774430 | 7.22 | -7.7 | -6.9 |
| CHEMBL3596868 | 6.92 | -9.0 | -6.2 |
| CHEMBL3596862 | 6.85 | -8.2 | -7.2 |
| CHEMBL1807198 | 6.72 | -8.8 | -6.5 |
| CHEMBL1170760 | 4.89 | -3.9 | -8.4 |
| CHEMBL3128218 | 4.82 | -7.2 | -6.6 |
| CHEMBL3329396 | 4.8 | -9.0 | -5.9 |
| CHEMBL4776658 | 4.79 | -7.1 | -6.6 |
| CHEMBL3128231 | 4.77 | -7.8 | -7.2 |
| CHEMBL3329401 | 4.76 | -6.8 | -5.6 |
| CHEMBL3329395 | 4.73 | -8.6 | -6.9 |
| CHEMBL4779979 | 4.72 | -8.6 | -7.3 |
| CHEMBL3329400 | 4.71 | na | -6.4 |
| CHEMBL601871 | 4.7 | -6.7 | -6.8 |
| CHEMBL3329399 | 4.69 | -1.5 | -5.5 |
| CHEMBL273103 | 4.68 | -4.0 | -4.3 |
| CHEMBL3128234 | 4.65 | -6.5 | -6.3 |
| CHEMBL3128232 | 4.62 | -1.4 | -5.8 |
| CHEMBL5193272 | 4.6 | -7.0 | -5.8 |
| CHEMBL1171688 | 4.58 | -7.9 | -7.7 |
| CHEMBL1170550 | 4.57 | -6.9 | -7.2 |
| CHEMBL3128233 | 4.55 | -8.2 | -7.1 |
| CHEMBL4450404 | 4.54 | -8.0 | -8.0 |

| | | | |
|---|---|---|---|
| CHEMBL4778459 | 4.5 | -7.8 | -7.2 |
| CHEMBL1704021 | 4.08 | -7.3 | -6.2 |
| CHEMBL172517 | 4.17 | -8.1 | -7.1 |
| CHEMBL1807517 | 4.26 | -8.8 | -7.6 |
| CHEMBL1807518 | 4.00 | -8.8 | -7 |
| CHEMBL1807519 | 4.00 | -8.1 | -6.5 |
| CHEMBL1807520 | 4.00 | -8.8 | -6.5 |
| CHEMBL1808239 | 4.00 | -8.1 | -8.2 |
| CHEMBL213207 | 4.00 | -8.2 | -7 |
| CHEMBL220444 | 4.00 | -6.5 | -4.2 |
| CHEMBL223486 | 4.00 | -7.0 | -7.4 |
| CHEMBL233373 | 4.18 | -8.2 | -5.4 |
| CHEMBL3128224 | 4.24 | -7.5 | -4.9 |
| CHEMBL3128226 | 4.23 | -7.6 | -6.7 |
| CHEMBL377048 | 4.00 | -8.6 | -6.8 |
| CHEMBL4237913 | 4.27 | -6.7 | -3.6 |
| CHEMBL4239930 | 4.20 | -6.9 | -6.3 |
| CHEMBL4537023 | 4.20 | -8.2 | -7.1 |
| CHEMBL4790509 | 4.32 | -5.0 | -4.9 |
| CHEMBL50 | 4.11 | -8.0 | -7.4 |
| CHEMBL597176 | 4.18 | -8.4 | -7.6 |
| CHEMBL597366 | 4.35 | -8.3 | -8.9 |
| CHEMBL603491 | 4.11 | -9.3 | -7.6 |
| CHEMBL608424 | 4.20 | -7.2 | -6.5 |
| CHEMBL90277 | 4.00 | -7.7 | -9.2 |
| CHEMBL213277 | 5 | -8.5 | -6.6 |

**Table S3.** Docking scores of specific PARP1 ligands using the ROC curve. Vina GPU 2.1 and DRGSCROLL were used to dock representative active chemicals with experimentally published pIC50 values.

| Name | $pIC_{50}$ | Vina GPU 2.1 (kcal/mol) | DRGSCROLL (kcal/mol) |
|---|---|---|---|
| CHEMBL3907010 | 9.7 | -8.3 | -7.9 |
| CHEMBL521686 | 9.39 | -9.7 | -9.6 |
| CHEMBL5395193 | 8.52 | -6.9 | -7.1 |
| CHEMBL5398113 | 8.52 | -8.4 | -8.2 |
| CHEMBL5408653 | 8.4 | -8.2 | -8.2 |
| CHEMBL5176058 | 8.38 | -10.3 | -10.3 |
| CHEMBL5196674 | 8.33 | -10.7 | -9.5 |
| CHEMBL5398856 | 8.3 | -6.7 | -7.6 |
| CHEMBL5404994 | 8.3 | -7.6 | -7.9 |
| CHEMBL5422478 | 8.3 | -7.1 | -8.8 |
| CHEMBL3644564 | 8.22 | -10 | -10.1 |
| CHEMBL3978133 | 8.22 | -9.5 | -8.5 |
| CHEMBL3936349 | 8.15 | -9.8 | -8.2 |
| CHEMBL5185911 | 8.09 | -10.3 | -8 |
| CHEMBL3644565 | 8.05 | -0.3 | -7.6 |
| CHEMBL3644616 | 8 | -5.4 | -7.3 |
| CHEMBL3644566 | 7.92 | -10.1 | -9.3 |
| CHEMBL3644605 | 7.89 | -5.7 | -9.1 |
| CHEMBL3644601 | 7.85 | -7.6 | -8.7 |
| CHEMBL3644598 | 7.82 | -7.1 | -9.1 |

| | | | |
|---|---|---|---|
| CHEMBL3644571 | 7.77 | -10.2 | -9.6 |
| CHEMBL3644604 | 7.77 | -7.2 | -8.1 |
| CHEMBL4169012 | 10.7 | -4.9 | -8.9 |
| CHEMBL5176160 | 9.39 | -9.4 | -8.0 |
| CHEMBL506871 | 10.21 | -9 | -7.9 |
| CHEMBL4170665 | 10 | -8.8 | -8.2 |
| CHEMBL5425729 | 9.77 | -1.2 | -9.3 |
| CHEMBL5403789 | 9.76 | -8.9 | -9.0 |
| CHEMBL4845834 | 9.72 | -4.3 | -10.5 |
| CHEMBL4159834 | 9.7 | -8.7 | -8.2 |
| CHEMBL4867855 | 9.64 | -7.7 | -9.1 |
| CHEMBL5428376 | 9.62 | -9.6 | -9.0 |
| CHEMBL5425393 | 9.59 | -10 | -9.3 |
| CHEMBL5095220 | 9.51 | -8.4 | -9.1 |
| CHEMBL5180001 | 9.47 | 6.2 | -6.0 |
| CHEMBL4171279 | 9.4 | 1.7 | -8.0 |
| CHEMBL5182785 | 9.39 | 6.7 | -9.2 |
| CHEMBL5395197 | 9.36 | -8.2 | -10.7 |
| CHEMBL4870960 | 9.33 | -4 | -8.6 |
| CHEMBL595679 | 9.3 | -2.6 | -7.9 |
| CHEMBL5089136 | 9.29 | -8.9 | -8.0 |
| CHEMBL3137320 | 9.24 | -8.9 | -9.6 |
| CHEMBL5193674 | 9.23 | 1.7 | -6.6 |
| CHEMBL4787850 | 9.22 | -7 | -8.7 |
| CHEMBL5208255 | 9.21 | -5.2 | -8.3 |
| CHEMBL3927445 | 7.57 | -9.3 | -9.6 |
| CHEMBL3644615 | 7.52 | -10 | -8.6 |
| CHEMBL2237224 | 7.38 | -8.5 | -10.4 |
| CHEMBL395610 | 7.25 | -8.2 | -7.7 |
| CHEMBL3290490 | 7.1 | -7.3 | -9.1 |
| CHEMBL4855558 | 7.3 | -9.2 | -10.2 |
| CHEMBL4165488 | 7 | -1.8 | -8.8 |
| CHEMBL5437680 | 6.98 | 12.1 | -8.0 |
| CHEMBL5434068 | 6.97 | -6.7 | -4.0 |
| CHEMBL5404678 | 6.85 | 25.7 | -6.6 |
| CHEMBL3987175 | 4.71 | -9.4 | -6.7 |
| CHEMBL3940683 | 4.71 | -7.7 | -6.2 |
| CHEMBL3986804 | 4.71 | -5.8 | -7.2 |
| CHEMBL3964642 | 4.71 | -8.9 | -7.6 |
| CHEMBL2419706 | 4.71 | -8.8 | -9.9 |
| CHEMBL3921853 | 4.7 | -8.4 | -9.8 |
| CHEMBL3918110 | 4.62 | -5.8 | -8.8 |
| CHEMBL3927023 | 4.53 | -8.6 | -7.8 |
| CHEMBL3902386 | 4.49 | -10.0 | -9.6 |
| CHEMBL2419716 | 4.15 | -9.7 | -9.5 |
| CHEMBL3941313 | 4.71 | -9.2 | -7.1 |
| CHEMBL3945237 | 4.71 | -7.6 | -10.2 |
| CHEMBL3945618 | 4.71 | -7.3 | -8.8 |
| CHEMBL3946798 | 4.71 | -8.4 | -6.5 |
| CHEMBL3956490 | 4.71 | -6.3 | -5.8 |
| CHEMBL3963547 | 4.37 | -9.1 | -8 |
| CHEMBL3964642 | 4.71 | -8.7 | -7.6 |
| CHEMBL3965278 | 4.71 | -8.6 | -9 |

| | | | |
|---|---|---|---|
| CHEMBL3975140 | 4.71 | -8.4 | -7.1 |
| CHEMBL3983088 | 4.71 | -8.5 | -7.9 |
| CHEMBL3985156 | 4.71 | -6.0 | -9.9 |
| CHEMBL4110589 | 4.71 | -9.3 | -8.4 |
| CHEMBL4111286 | 4.71 | -6.8 | -9.9 |
| CHEMBL4112739 | 4.71 | -9.0 | -7.9 |
| CHEMBL4113551 | 4.71 | -6.1 | -4.7 |
| CHEMBL3416132 | 4 | -7.7 | -7.0 |
| CHEMBL1688212 | 4 | -7.3 | -6.9 |
| CHEMBL2323227 | 4.06 | -10 | -9.5 |
| CHEMBL2323532 | 4.08 | -11 | -8.6 |
| CHEMBL1813880 | 4.11 | -6 | -9.4 |
| CHEMBL2323531 | 4.12 | -8 | -10.4 |
| CHEMBL4099515 | 4.13 | -10 | -9.9 |
| CHEMBL1813887 | 4.18 | 9.1 | -6.8 |
| CHEMBL5433754 | 4.2 | -6.8 | -6.6 |
| CHEMBL480233 | 4.21 | -7.6 | -8.8 |
| CHEMBL4797518 | 4.23 | -7.1 | -7.8 |
| CHEMBL449635 | 4.24 | -6.4 | -5.5 |
| CHEMBL122690 | 4.26 | -6.6 | -6.4 |
| CHEMBL479635 | 4.28 | -8.7 | -8.8 |
| CHEMBL5271762 | 4.31 | -9.2 | -7.9 |
| CHEMBL4069154 | 4.86 | -10.3 | -9.5 |
| CHEMBL480427 | 4.37 | -8 | -8.9 |
| CHEMBL3929392 | 4.43 | -4.6 | -9.6 |
| CHEMBL1086580 | 4.46 | -6.9 | -8.3 |
| CHEMBL1767055 | 4.5 | -9.3 | -9.0 |
| CHEMBL123978 | 4.52 | -6.5 | -5.7 |
| CHEMBL3907010 | 9.7 | -8.3 | -7.9 |

**Table S4.** Docking scores of active and decoys ligands selected from DUDE for PARP1 using the ROC curve. Vina GPU 2.1 and DRGSCROLL were used to dock representative active chemicals.

| Name | DRGSCROLL (kcal/mol) | Vina GPU 2.1 (kcal/mol) |
|---|---|---|
| CHEMBL108339 | -9.4 | -5.1 |
| CHEMBL1088788 | -11 | 11.1 |
| CHEMBL1094951 | -10.8 | -9.2 |
| CHEMBL110729 | -12 | -8.6 |
| CHEMBL188985 | -10.1 | 12.4 |
| CHEMBL193918 | -13.4 | -9.7 |
| CHEMBL194416 | -7.3 | -5.6 |
| CHEMBL196450 | -14.2 | -10.2 |
| CHEMBL320579 | -11.7 | -8.3 |
| CHEMBL338955 | -10.9 | -8.7 |
| CHEMBL383442 | -14.8 | -10.3 |
| CHEMBL432211 | -12.8 | -10.4 |
| CHEMBL471966 | -10 | -8.1 |

| | | |
|---|---|---|
| CHEMBL516910 | -11.3 | -4.7 |
| CHEMBL520745 | -11.9 | -10.3 |
| CHEMBL595230 | -3.2 | 67.8 |
| CHEMBL595924 | -3.6 | 7.3 |
| CHEMBL596144 | -12.4 | -8.9 |
| CHEMBL597541 | -13.8 | -8.3 |
| CHEMBL611980 | -10.5 | -5.8 |
| C00048594 | -10.5 | -9 |
| C00174747 | -10.4 | -8.4 |
| C00195967 | -11.6 | -8.9 |
| C00371786 | -9.5 | -8 |
| C00612924 | -10.6 | -8.7 |
| C02107078 | -10.4 | -4.4 |
| C02484133 | -10 | -8.2 |
| C02784705 | -8.5 | -4.8 |
| C02798507 | -6.2 | -5.1 |
| C03847560 | -11.9 | -9.2 |
| C04228176 | -8.1 | -6.5 |
| C04682082 | -10.8 | -9.7 |
| C04909264 | -11.5 | -9.5 |
| C04945114 | -10.5 | -3.8 |
| C05145970 | -12 | -10 |
| C05176737 | -10.8 | -1.1 |
| C05721576 | -8.1 | -6.7 |
| C60208788 | -12.3 | -7.3 |
| C61846715 | -9.3 | -7.9 |
| C61886223 | -9.7 | -7.9 |
| C62496970 | -9.7 | -7.1 |
| C62638937 | -11.5 | -9.5 |
| C63039540 | -7.6 | -6.5 |
| C63039549 | -7 | -5.9 |
| C63513821 | -9.9 | -8.2 |
| C63853760 | -9.7 | -6.7 |
| C64821621 | -8.8 | 33.8 |
| C64965926 | -9.7 | 3.7 |
| C65337591 | -7.1 | -5.8 |
| C65407009 | -12.3 | -8.4 |
| C66239617 | -7.9 | 23.8 |
| C66239618 | -9.2 | 25.8 |
| C66239705 | -9 | 12 |
| C66240207 | -7.9 | 13.9 |
| C66241463 | -11.4 | 18.8 |
| C66246208 | -9.1 | -8 |

| | | |
|---|---|---|
| C66249629 | -11.4 | -7.3 |
| C66286865 | -10.5 | -8.5 |
| C66402266 | -10 | -7.9 |
| C66708893 | -12.5 | -8.8 |
| C66710978 | -12.5 | -8.8 |
| C66741192 | -11.7 | -9.2 |
| C66783620 | -9.2 | 2.5 |
| C66897726 | -9 | 0.1 |
| C66908495 | -11.6 | -8.9 |
| C66913869 | -9.6 | -7.9 |
| C67000332 | -10 | -5.3 |
| C67051502 | -10 | -7.9 |
| C67237465 | -11.7 | -7.3 |
| C67386459 | -11.1 | -8.7 |
| C06389686 | -10.4 | -6.1 |
| C40143378 | -11.8 | -8.7 |
| C40359701 | -10 | -7.9 |
| C40570637 | -11.4 | -8.8 |
| C40921264 | -13 | -7.8 |
| C41021313 | -9.7 | -7.7 |
| C41249834 | -9.2 | -7.8 |
| C42006984 | -11.1 | -8.8 |
| C42381385 | -11.7 | -9.6 |
| C42610406 | -10.1 | -8.2 |
| C42821586 | -11.2 | -8.3 |
| C43218608 | -10.3 | -5.5 |
| C43741785 | -10.2 | -6.7 |
| C43942831 | -11.8 | -7.5 |
| C43942832 | -11.4 | -7.9 |
| C44449716 | -10.5 | -7.9 |
| C44783086 | -9.7 | -8.1 |
| C44944297 | -10.5 | -8.2 |
| C45147868 | -9 | -7.4 |
| C45274685 | -11.3 | -8.6 |
| C45628394 | -7.4 | -6.6 |
| C45644043 | -10.6 | -2.5 |
| C45827843 | -9.3 | -5.8 |
| C47553589 | -9.9 | -2.7 |
| C48045922 | -9.2 | -7.6 |
| C48278950 | -8.5 | -3.8 |
| C49060597 | -9.6 | 10.5 |
| C49524243 | -11.5 | -3.8 |
| C49587745 | -8.3 | -6.9 |

| | | |
|---|---|---|
| C50002172 | -10.2 | -7.8 |
| C50842985 | -10.1 | -7.4 |
| C50892214 | -10.4 | -8.3 |
| C51114300 | -9.1 | -7.5 |
| C51269002 | -10.4 | -5.7 |
| C51687658 | -8.5 | -7.1 |
| C52068402 | -9.4 | -7.8 |
| C53756474 | -10.6 | -7.4 |
| C54118371 | -8.9 | -5.7 |
| C54533024 | -10.2 | -8.4 |
| C55053194 | -11.9 | -9.4 |
| C55266209 | -10.2 | -7.9 |
| C55399946 | -12.1 | -7.7 |
| C56748541 | -10.8 | -7.4 |
| C56937284 | -9.9 | -8.3 |
| C57568064 | -9.5 | -8.3 |
| C58245025 | -8.7 | 22.4 |
| C58329780 | -9.7 | -7.5 |
| C58425222 | -7.6 | -6 |
| C59327058 | -11.2 | -9.1 |
| C06831244 | -11.7 | -7.7 |
| C07077896 | -10 | -7.7 |
| C07424297 | -11.6 | -8.1 |
| C08594938 | -8.7 | -2.2 |
| C09118156 | -11.9 | -5.1 |
| C09409795 | -10 | -7.3 |
| C09439708 | -10.7 | -8.6 |
| C09536376 | -11.6 | -9.8 |
| C09600346 | -11.2 | -8.3 |
| C09874757 | -10.2 | -8.3 |
| C10241719 | -9.1 | -6.8 |
| C10895098 | -9.4 | 6.9 |
| C11904448 | -10.4 | -8.8 |
| C11976484 | -10.1 | 44.2 |
| C11984102 | -9 | -7.5 |
| C12031574 | -7 | -5.7 |
| C12275366 | -12.3 | -8.7 |
| C12316179 | -11.4 | 8 |
| C12341480 | -9.3 | -6.7 |
| C12349864 | -13.2 | -9.3 |
| C12410457 | -7.4 | -5.5 |
| C12624103 | -11.9 | -8.5 |
| C12820352 | -10.6 | -8.2 |

| | | |
|---|---|---|
| C12859872 | -11.1 | -9.2 |
| C13101932 | -9 | -6.5 |
| C13165367 | -10.1 | -7.7 |
| C14016618 | -11.4 | -4.9 |
| C14071137 | -12.5 | -10.4 |
| C14110886 | -10.9 | -8.7 |
| C14186071 | -7.5 | -5.9 |
| C14530914 | -10.8 | -4.7 |
| C16260947 | -9 | -6.4 |
| C17378513 | -11.1 | -8.9 |
| C18066485 | -12 | 2.4 |
| C18080918 | -13.3 | -10 |
| C18136750 | -12.7 | -9.5 |
| C20131175 | -10.3 | -2.3 |
| C20149485 | -11.7 | -9.9 |
| C20327112 | -12.9 | -8.7 |
| C20392819 | -10.2 | 1.2 |
| C20745277 | -11.1 | -9.4 |
| C21706802 | -11.2 | -7.7 |
| C21866358 | -11.3 | -8.6 |
| C22057864 | -11.5 | -7.7 |
| C22693106 | -11 | -6.7 |
| C22872385 | -9.7 | 2.6 |
| C23158434 | -9.2 | 14.6 |
| C23334772 | -7.4 | 4.6 |
| C24960808 | -11.3 | -9.1 |
| C25400943 | -9.9 | 12 |
| C25601885 | -9.9 | -7.1 |
| C26442999 | -9.4 | -8.6 |
| C26578801 | -8.2 | 14.5 |
| C26828762 | -9.9 | -8.3 |
| C27222456 | -11.7 | -8.4 |
| C28560429 | -7.5 | 10.6 |
| C30058540 | -10.4 | -5.6 |
| C30550976 | -10.6 | -6.4 |
| C30816379 | -10.1 | -5.9 |
| C32070740 | -11.7 | -8 |
| C32536495 | -9.6 | -7.6 |
| C32606782 | -10.9 | 2.3 |
| C32883647 | -10.7 | -7.4 |
| C33121632 | -8.4 | 32 |
| C33419736 | -10.2 | -8.8 |
| C33812507 | -9.9 | -0.1 |

| | | |
|---|---|---|
| C33931134 | -12.3 | -7.2 |
| C34776612 | -9.6 | -6.7 |
| C34786022 | -10.3 | 1 |
| C34786301 | -8.3 | -1.6 |
| C35422361 | -10 | -7.3 |
| C35526635 | -10.4 | -8.5 |
| C35674576 | -9.3 | -6.9 |
| C36006190 | -11.4 | -2.7 |
| C36210478 | -8.1 | -0.5 |
| C36596542 | -10.1 | -7 |
| C36597181 | -9.8 | -7.3 |
| C36667012 | -8.7 | -7.2 |
| C36730974 | -9.9 | -6.7 |
| C36740558 | -12 | -9.2 |
| C36749549 | -8.7 | -7.5 |
| C36809138 | -9.3 | -7 |
| C36929090 | -8.1 | -7 |
| C36938926 | -8.7 | -7.4 |
| C37641946 | -10.7 | -8.3 |
| C37694578 | -7.6 | -5.7 |
| C37755194 | -7 | -5.7 |
| C37859412 | -10.4 | -8.3 |
| C38685202 | -10.3 | -8.6 |
| C38695428 | -11.7 | -6 |
| C38695444 | -11.7 | -6 |
| C39104593 | -6.8 | -5.1 |
| C39161249 | -10.8 | -8.8 |
| C39354102 | -11.3 | -6.7 |
| C39440133 | -8.7 | -7 |
| C39855129 | -11.8 | -8.2 |
| C39958083 | -13 | -9.1 |
| C39958085 | -13.7 | -10.6 |
| C40021225 | -11.1 | -5.5 |
| C58224937 | -9.4 | 8.3 |
| C06441966 | -10.5 | -8.9 |

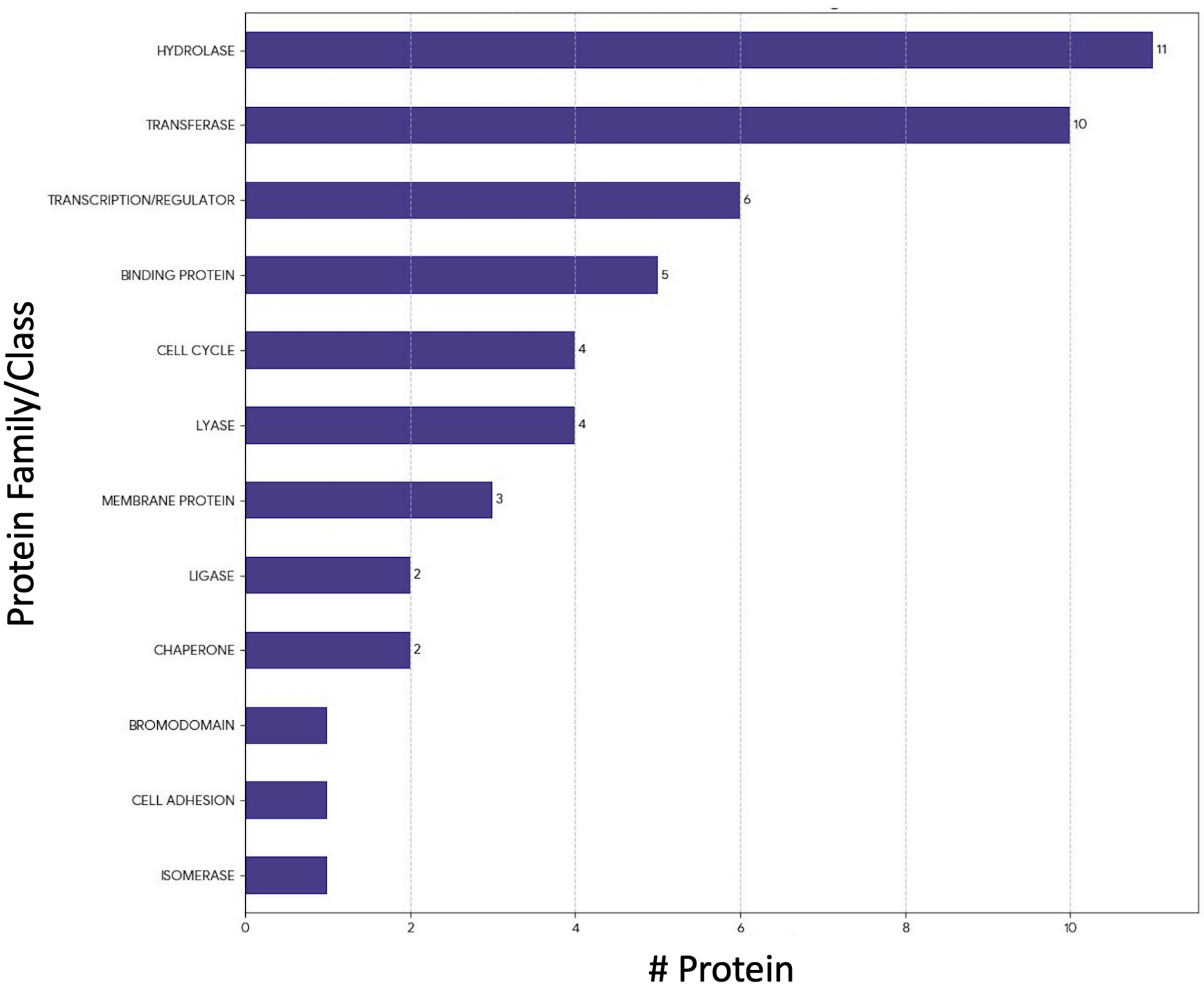


**Figure S1.** Random selection of 50 protein-ligand complexes. Figure represents number of protein families per protein family.

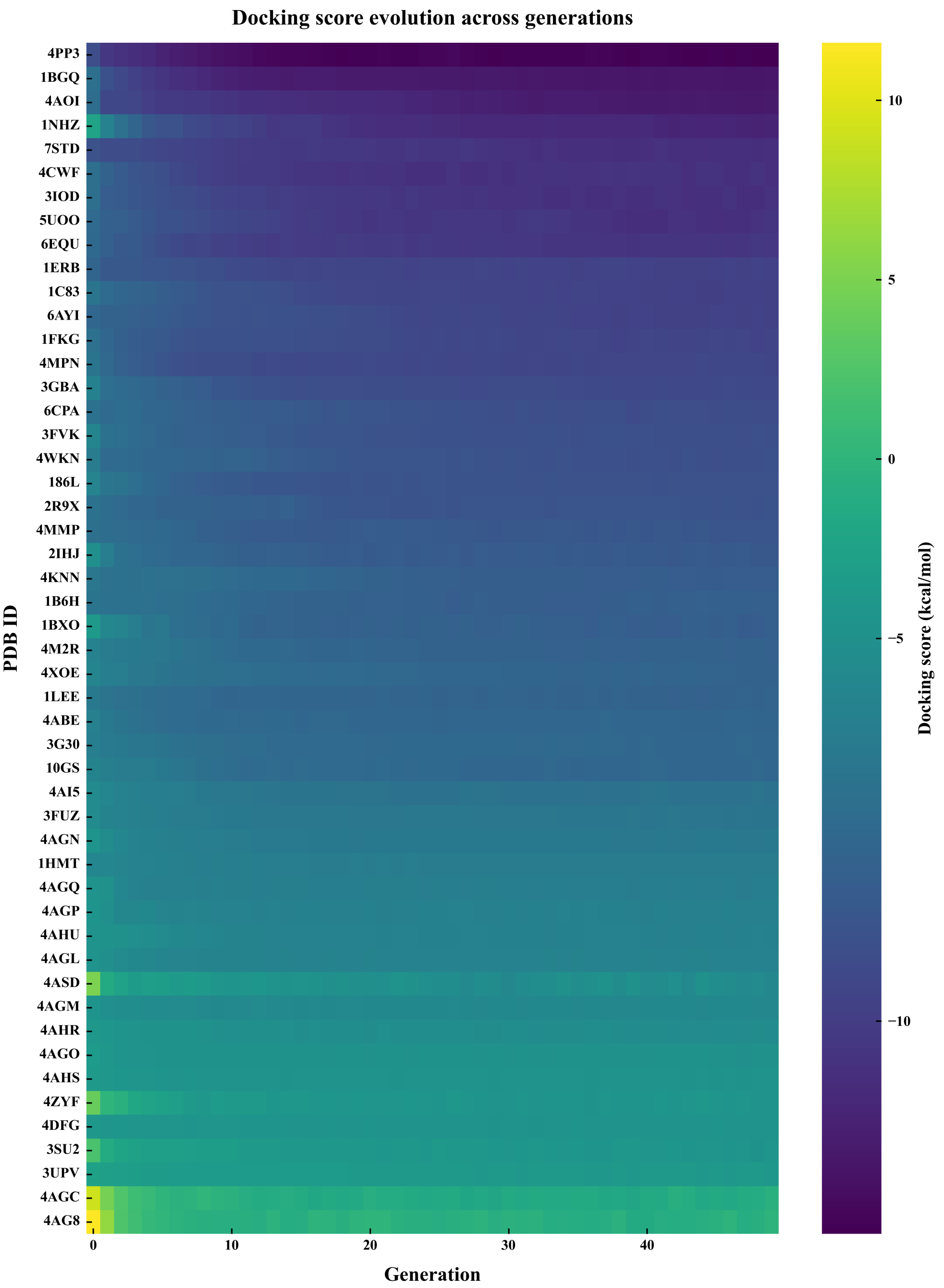


**Figure S2.** Docking score evolution for 50 protein targets used throughout generations. Within the GA-based flexible docking workflow, each column denotes a generation, and each row represents a unique protein target (PDB ID). Each generation's docking scores were averaged among population members. Stronger anticipated binding affinities are indicated by lower (purple–blue) values on the viridis color scale, which is centered at –5 kcal $mol^{-1}$. Weaker or unfavorable postures are indicated by higher (yellow–green) values. According to the heatmap, most systems gradually move toward lower docking energies over the course of subsequent generations, suggesting evolutionary convergence toward ligand–protein complexes that are energetically advantageous.

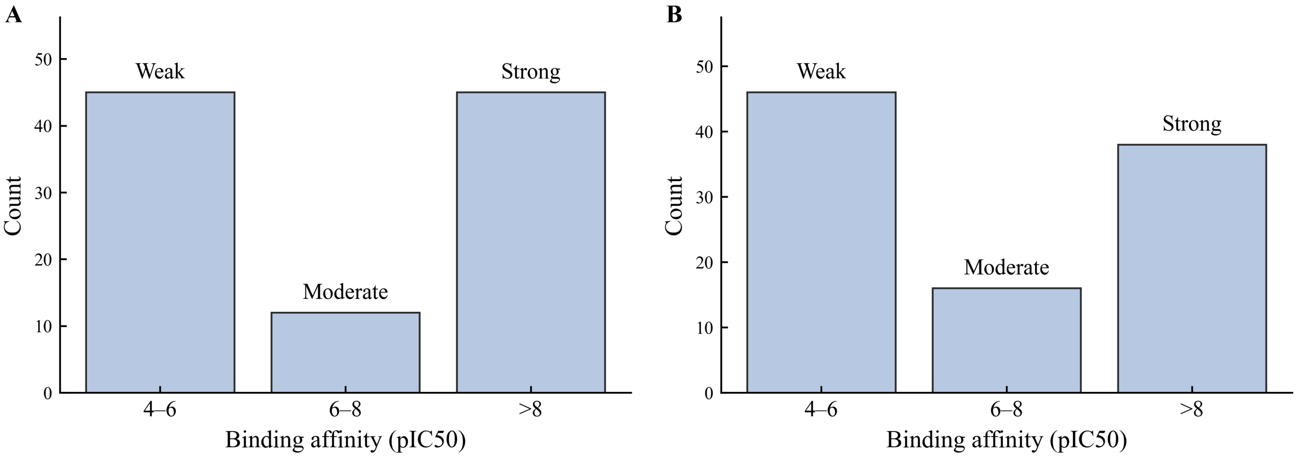


**Figure S3.** Compound distribution according to types of binding affinity. (A–B) Three affinity ranges were identified based on the experimental $pIC_{50}$ values of the compounds: mild ($4 \leq pIC_{50} < 6$), moderate ($6 \leq pIC_{50} < 8$), and strong ($pIC_{50} \geq 8$). The amount of compounds in each category is shown by bar heights. The matching affinity class is shown by labels above the bars, and the $pIC_{50}$ range bounds are shown on the x-axis. The majority of molecules in both datasets fall into the weak and strong binding ranges, whereas the moderate group is underrepresented, indicating a bimodal distribution of ligand potencies.

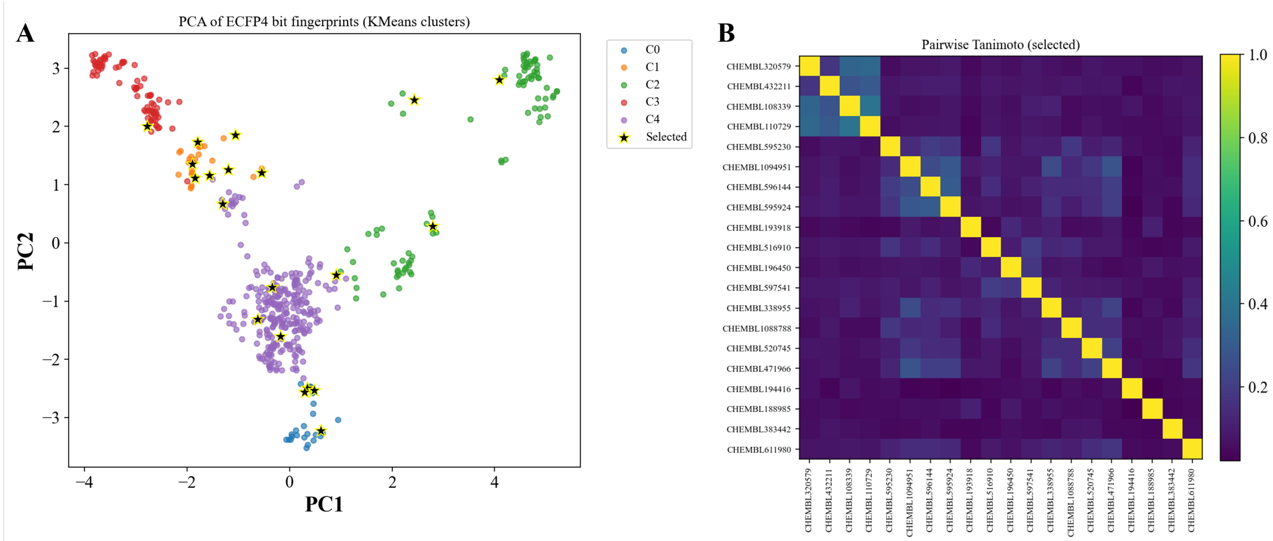


**Figure S4.** DUDE-PARP1 active compound diversity evaluation and cluster-based selection. (A) KMeans (k=5) clustered ECFP4 (radius 4, 4096-bit) molecular fingerprints visualized using PCA. The 20 chosen sample molecules are indicated by yellow stars, and each point represents an active chemical colored by its allocated cluster. (B) A pairwise Tanimoto similarity heatmap of the 20 actives that were chosen reveals a significant level of chemical diversity among the representatives and a low level of overall similarity.

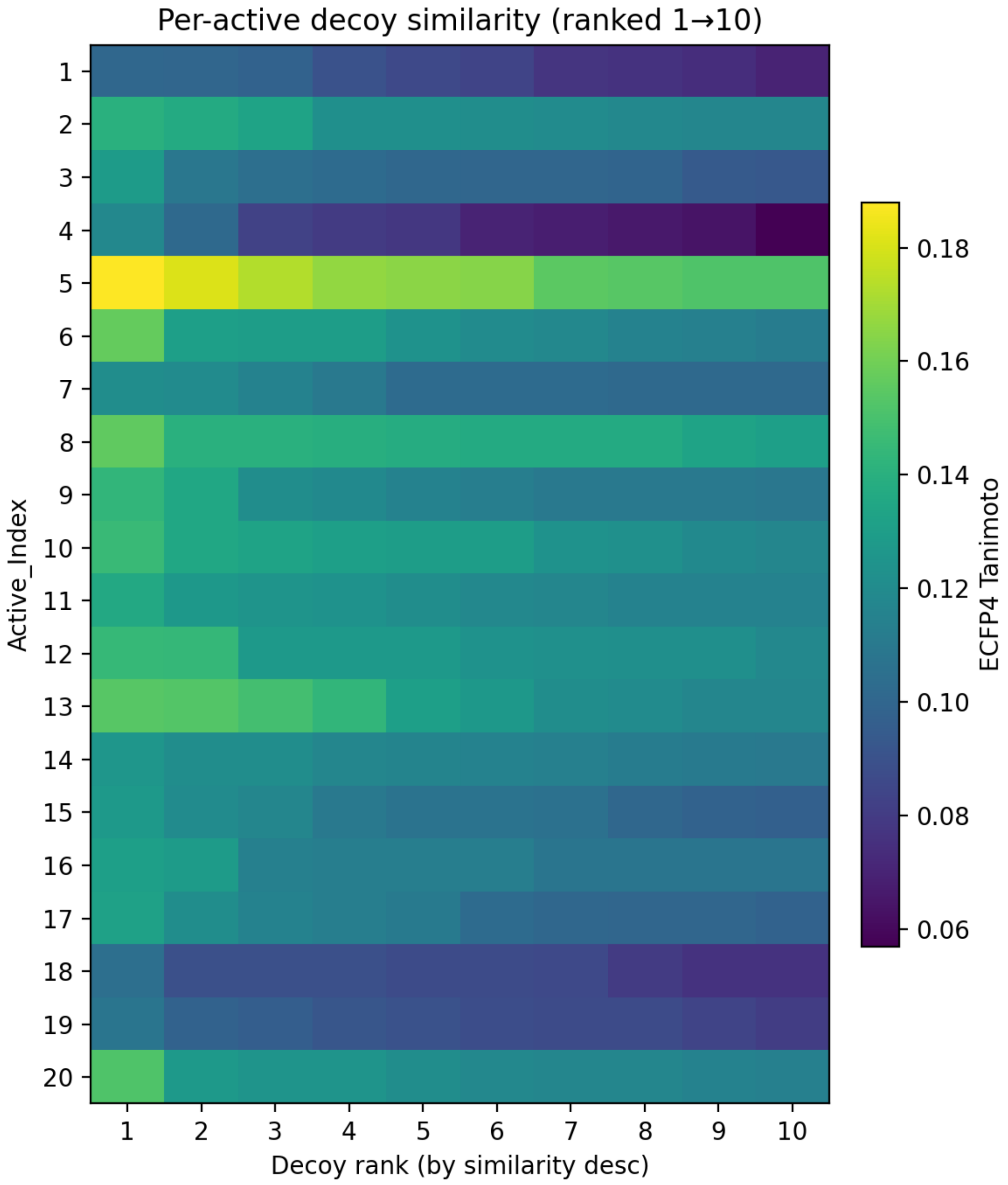


**Figure S5.** ECFP4 Tanimoto values per-active heatmap (rows: actives 1–20; columns: their 10 decoys ranked by decreasing similarity). With typical ranges of 0.10 to 0.18 and a concentration around T = 0.13, the similarity distributions confirm little structural overlap while maintaining physicochemical matching.

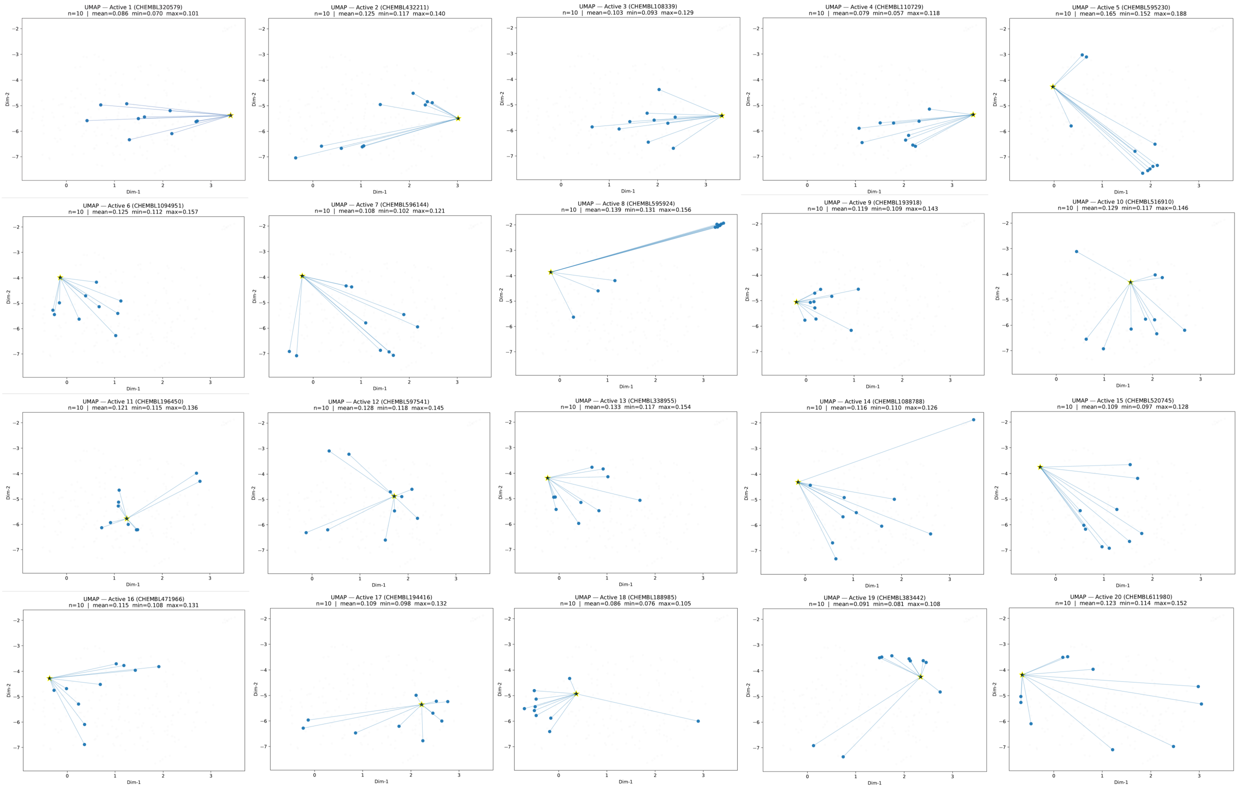


**Figure S6.** Each active star (yellow) and its ten decoys (blue) are projected by UMAP. Decoys show chemically plausible but structurally different negatives that are appropriate for virtual-screening benchmarks by populating proximal regions without collapsing over the active.
